# Chemical factors induce aggregative multicellularity in a close unicellular relative of animals

**DOI:** 10.1101/2022.12.01.517680

**Authors:** Núria Ros-Rocher, Ria Q. Kidner, Catherine Gerdt, W. Sean Davidson, Iñaki Ruiz-Trillo, Joseph P. Gerdt

## Abstract

Regulated cellular aggregation is an essential process for development and healing in many animal tissues. In some animals and a few distantly related unicellular species, cellular aggregation is regulated by diffusible chemical cues. However, it is unclear whether regulated cellular aggregation was part of the life cycles of the first multicellular animals and/or their unicellular ancestors. To fill this gap, we investigated the triggers of cellular aggregation in one of animals’ closest unicellular living relatives – the filasterean *Capsaspora owczarzaki*. We discovered that *Capsaspora* aggregation is induced by chemical cues, as observed in some of the earliest branching animals and other unicellular species. Specifically, we found that calcium ions and lipids present in lipoproteins function together to induce aggregation of viable *Capsaspora* cells. We also found that this multicellular stage is reversible, as depletion of the cues triggers disaggregation, which can be overcome upon re-induction. Our finding demonstrates that chemically regulated aggregation is important across diverse members of the holozoan clade. Therefore, this phenotype was plausibly integral to the life cycles of the unicellular ancestors of animals.

## INTRODUCTION

Cellular aggregation arises through the clustering and adhesion of individual cells. This multicellular behavior is broadly distributed across the eukaryotic tree of life, being particularly common in various unicellular organisms **(Fig. 1A)** (1–5). Many of these species, like the amoebozoan *D. discoideum*, utilize chemical cues to regulate aggregation in response to harsh environmental conditions (3, 6–16).

**Figure 1.**
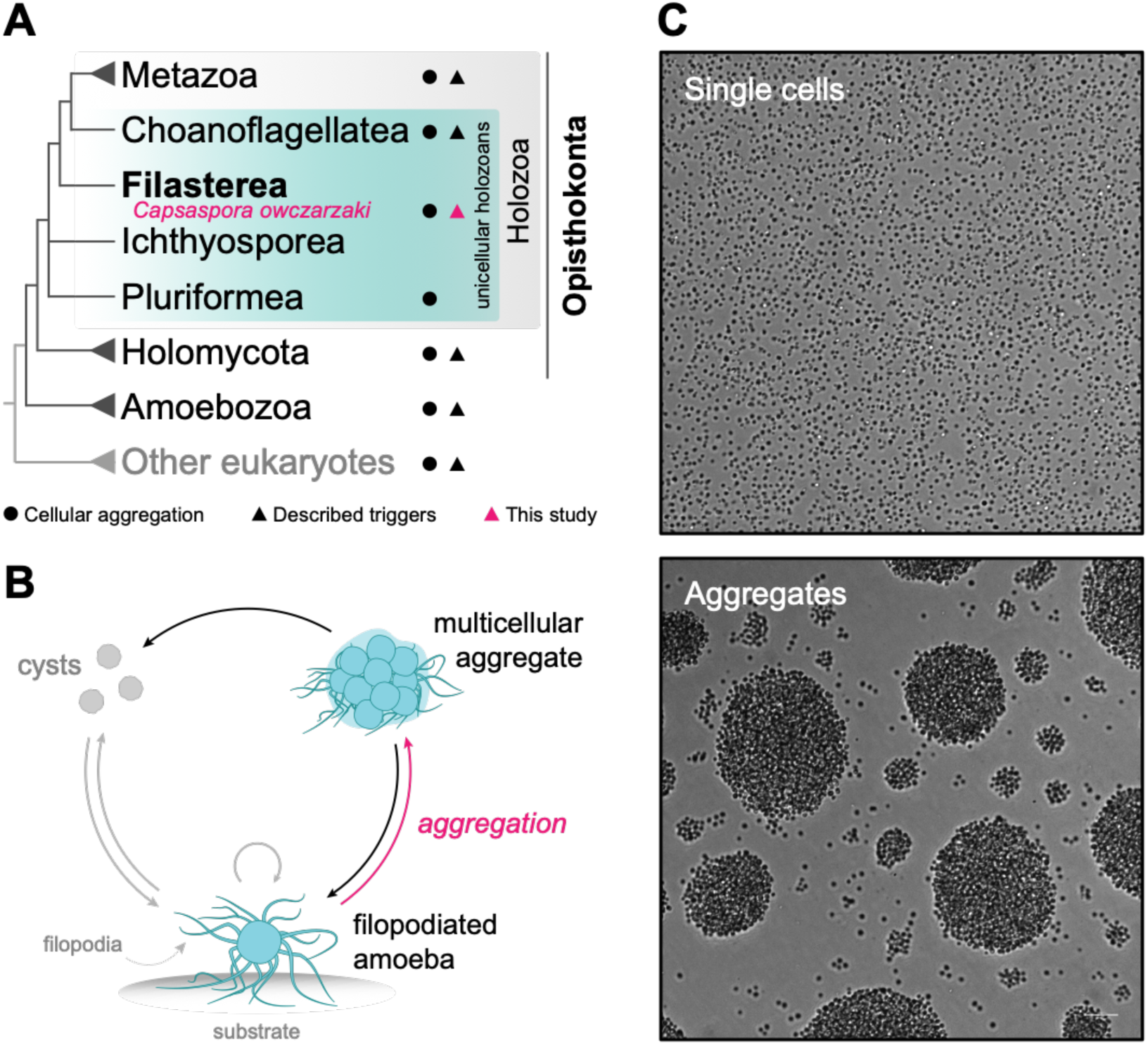
*Capsaspora owczarzaki* transitions from a unicellular to a multicellular life stage. (A) Animals (Metazoa) and their closest unicellular relatives, including *Capsaspora* (Filasterea), form the Holozoa clade within the Opisthokonta supergroup. The phylogenetic relationships of selected taxa are based on several recent phylogenomic studies (76–78). Uncertain positions are represented with polytomies. (B) Life stages of *Capsaspora* under culture conditions. The trophic proliferative cells are filopodiated amoebae adhered to the substrate (adherent stage). These amoebae can detach from the substrate and form multicellular aggregates (aggregative stage). In response to crowding or stress, cells from both the adherent and the aggregative stages can encyst by retracting the filopodia (cystic stage). Arrows indicate directionality of each transition, with an emphasis on aggregation (pink arrow) and disaggregation (black arrows). Loop arrow indicates cell division in adherent cells (35, 58). (C) *Capsaspora* single cells at the adherent stage (top panel) transition to multicellular aggregates (bottom panel). Scale bar represents 100 µm.

Diverse taxa of animals (Opisthokonta, Holozoa) **(Fig. 1A)** also exhibit cellular aggregation during fundamental processes to develop and maintain healthy multicellular tissues. From sponges and cnidarians to vertebrates, animal cells disaggregate and re-aggregate for spatial organization of tissues during the early stages of embryonic development (17–19), during inflammatory processes (20, 21), and for tissue regeneration (22–29). A few animal species have been conclusively demonstrated to use soluble chemical cues to regulate their aggregation phenotypes (6, 23, 30, 31). However, it is unclear how chemically regulated cellular aggregation evolved in the animal linage and whether it was part of the life cycles of the latest unicellular ancestors of animals.

Recently, extant unicellular relatives of animals (*i.e.,* unicellular holozoans, **Fig. 1A**) have become powerful models to reconstruct which genes and biological traits were likely present in the unicellular ancestors of animals (32, 33). For instance, comparative genomic studies between animals and unicellular holozoans revealed that the genomes of many unicellular holozoans encode genes homologous to those that are essential for animal development and multicellularity. These include components of the animal cell adhesion machinery, developmental transcription factors, and cell-cell communication proteins, which are thus inferred to be present in their common unicellular ancestors (32, 34). Similarly, analyses of their life styles showed that unicellular holozoans exhibit a diversity of multicellular phenotypes, including multicellular colonies formed through cellular aggregation (35, 36). These findings suggest that the life cycles of the unicellular ancestors most likely integrated single-celled states and transient multicellular phenotypes. However, it remains obscure how unicellular ancestors regulated most multicellular phenotypes, and particularly, whether they could have employed similar mechanisms to those observed in animals.

Among unicellular holozoans, the filasterean amoeba *Capsaspora owczarzaki,* hereafter *Capsaspora,* can form multicellular aggregates (35, 37, 38) that are morphologically similar to aggregates formed by animal cells, particularly sponges and hydra during whole-body regeneration **(Fig. 1B-C)** (22, 26, 27, 35). These animal-like *Capsaspora* aggregates exhibit elevated expression of many genes homologous to those involved in cell adhesion in animals (35, 37, 38). Nevertheless, little is known about how *Capsaspora* regulates its aggregative phenotype.

Therefore, we sought to characterize how *Capsaspora* regulates cellular aggregation and compare it to aggregation regulation in present-day animals and other aggregating unicellular species. Because some animal cells and unicellular species form aggregates in response to chemical cues (7, 8, 22, 23, 39–42), we specifically asked if *Capsaspora* also employs extracellular chemical signals to regulate its aggregative behavior. We discovered that *Capsaspora* aggregation, like cellular aggregation in some animals, choanoflagellates, and some other unicellular species, indeed depends on chemical cues. We identified the chemical cues to be calcium ions and lipids present in lipoproteins. We also discovered that *Capsaspora* aggregation is a reversible process that depends on the concentration of the cues. Moreover, the cues only induce aggregation of viable *Capsaspora* cells, suggesting that an active cellular response regulates aggregation. Further molecular studies are needed to determine the degree to which this regulated aggregation behavior is genetically conserved with animals. Nonetheless, this finding adds to the increasing prevalence of chemically regulated multicellularity in unicellular animal relatives, which suggests that the close unicellular ancestors of animals may have been capable of chemically regulated multicellular phenotypes.

## RESULTS

### Fetal Bovine Serum contains chemical factor(s) necessary for aggregation in *Capsaspora*

*Capsaspora* aggregates have previously been induced using physical forces by gentle orbital agitation of amoeboid cells in a manner similar to sponge primmorph induction (22, 35, 37, 38). Since some animals and other unicellular species secrete chemical cues to promote cell aggregation (7, 41, 43, 44), we initially inquired whether *Capsaspora* aggregates could be induced by secreted cues from aggregating cultures under agitation conditions. However, preliminary experiments testing conditioned medium from aggregated cultures did not result in aggregation of amoeboid cells (data not shown).

We then asked whether environmental chemical factors could be necessary to induce aggregation in *Capsaspora*. During the aggregation process, *Capsaspora* cells are maintained in growth media: a nutrient-rich buffered solution containing 1% (w/v) peptone, 1% (w/v) yeast extract, and 10% (v/v) heat-inactivated Fetal Bovine Serum (FBS). To determine if any of these components were necessary for aggregation, we agitated *Capsaspora* cultures in variations of the media lacking each of these components. We first modified the previously reported aggregation assay (35) to test multiple conditions and reduce the amount of sample used **(Fig. S1A)** and also incorporated an automated image analysis pipeline to distinguish between aggregates and single cells based on particle size **(Fig. S2)**. Using this setup, we found that cells agitated in FBS-free media were unable to form aggregates, suggesting that chemical cues from FBS are required to induce aggregation in *Capsaspora* **(Fig. 2A-B)**.

**Figure 2.**
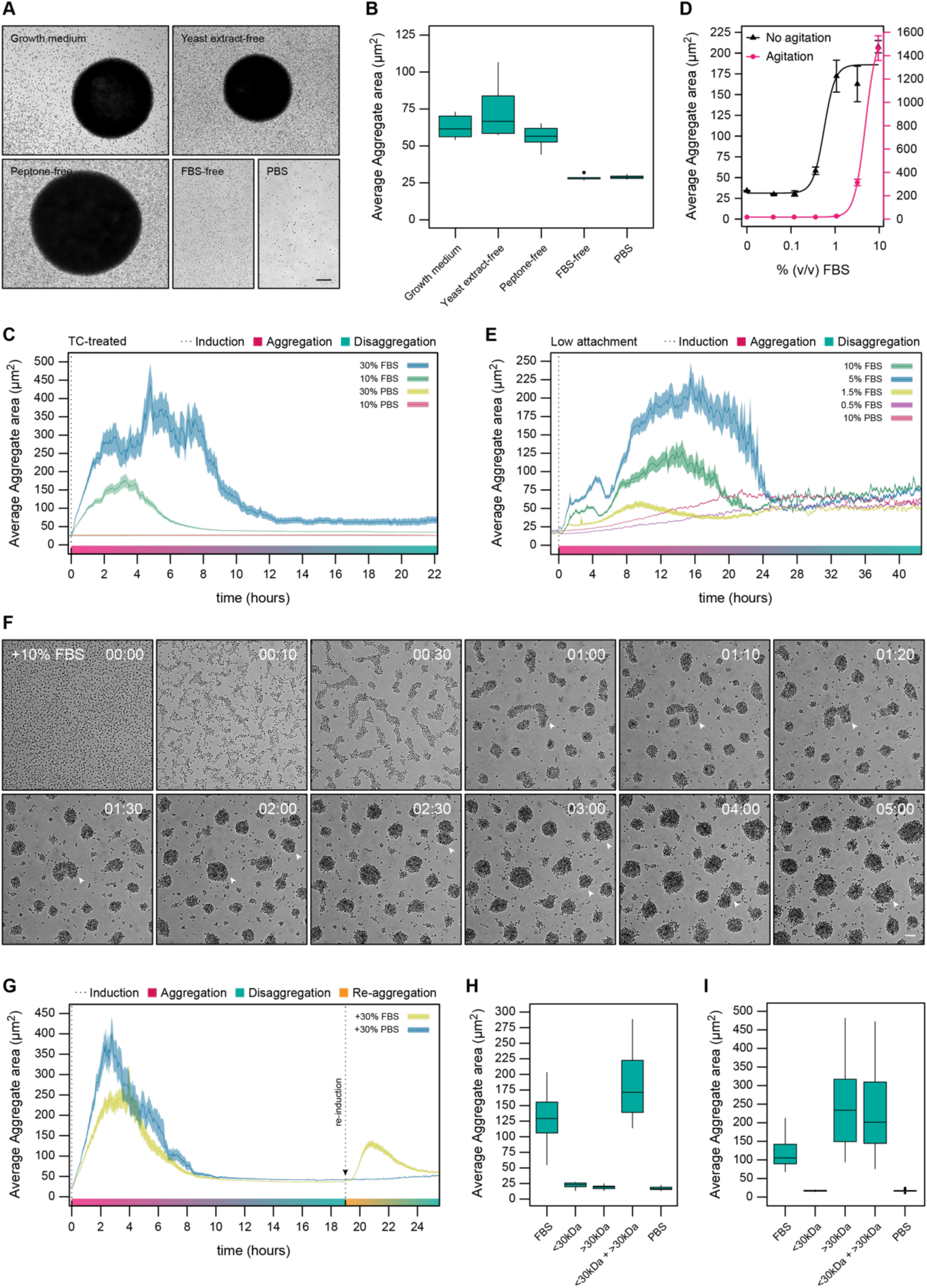
Chemical cues from FBS are necessary to induce aggregation in *Capsaspora.* (A) Microscopy images of *Capsaspora* aggregates induced under agitation conditions in growth medium and medium lacking some of its major components. Aggregates failed to form in FBS-free media and in 1X PBS (negative control) **(Fig. S1A assay)**. Scale bar represents 100 µm. (B) Data from each test condition in A are represented as boxplots, showing the median aggregate area (thick black bar) and interquartile ranges from three independent biological replicates. Figure related to **Fig. S2.** (C) FBS is sufficient to induce aggregation in the absence of physical agitation. Aggregation dynamics over time induced by 10% and 30% (v/v) FBS in the absence of physical agitation using tissue culture-treated (TC-treated) 12-well plates **(Fig. S1C assay)**. A 10% (v/v) and 30% (v/v) of 1X PBS were used as negative controls. Data are represented as the average aggregate area (thick line) ± s.e.m. (shadow) from three biological replicates. Quantification plot related to **Movie S1** and **Movie S2**. (D) Aggregation of *Capsaspora* cells induced by dose-responses of FBS in agitation **(Fig. S1B assay)** and in the absence of agitation conditions **(Fig. S1D assay)**. Data are represented as the average aggregate area ± s.e.m. from three independent biological replicates. (E) Aggregation dynamics over time induced by serial dilutions of FBS (v/v) in the absence of agitation conditions in ultra-low attachment 96-well plates **(Fig. S1D assay)**. A 10% (v/v) of 1X PBS was used as negative control. Data are represented as the average aggregate area (thick line) ± s.e.m. (shadow) from three independent biological replicates in no agitation conditions. Quantification plot related to **Movie S3** and **Movie S4**. (F) Aggregate formation induced by 10% (v/v) of FBS (time 0) in ultra-low attachment plates related to **Movie S3 (Fig. S1D assay and Fig. S3)**. White arrowheads track early aggregate formation and aggregate fusion events (agglomeration). Time represents hh:mm. Scale bar represents 100 µm. (G) Aggregation and re-aggregation dynamics over time in the absence of physical agitation using 12-well plates **(Fig. S1C assay)**. Aggregates were induced with a 10% (v/v) of FBS (time 0) and monitored until disaggregation into single-cells. After ∼19h, either a 30% (v/v) of FBS was added to induce re-aggregation (yellow curve) or a 30% (v/v) of 1X PBS was added as negative control (blue curve). Data are represented as the average aggregate area (thick line) ± s.e.m. (shadow) from three biological replicates. Quantification plot related to **Movie S5**. A similar result was observed for re-aggregation induced with 10% (v/v) of FBS **(Fig. S4)**. (H) The combination of <30 kDa and >30 kDa FBS fractions induce aggregation of *Capsaspora* cells. Cells were induced with 10% (v/v) of whole FBS, 10% (v/v) of <30 kDa FBS fraction, 10% (v/v) of >30 kDa FBS fraction or a combination of both fractions (5% (v/v) each) in agitation conditions **(Fig. S1B assay).** A 10% (v/v) of 1X PBS was used as a negative control. Data are represented as boxplots, showing the median aggregate area (thick black bar) and interquartile ranges from three independent biological replicates. (I) The >30 kDa FBS fraction is sufficient to induce re-aggregation of *Capsaspora* cells. Aggregation was induced with 10% (v/v) of FBS and monitored until disaggregation (∼72h). At this point, disaggregated cells were re-induced with either 10% (v/v) of FBS, 10% (v/v) of <30 kDa FBS fraction, 10% (v/v) of >30 kDa FBS fraction or a combination of both fractions (5% (v/v) each) in agitation conditions **(Fig. S1B assay).** A 10% (v/v) of 1X PBS was used as a negative control. Data are represented as boxplots, showing the median aggregate area (thick black bar) and interquartile ranges from three independent biological replicates.

### Chemical induction is sufficient for aggregation, even in the absence of physical agitation

In the process of assessing the aggregation effect of FBS, we serendipitously observed that *Capsaspora* cells can aggregate without physical agitation. In fact, cells in FBS-free media formed small aggregates within 20 minutes after addition of 10% (v/v) FBS *in the absence of orbital agitation* **(Fig. 2C, Fig. S1C, Movie S1)**. This aggregation effect was even more pronounced upon addition of 30% (v/v) FBS, which induced larger aggregates within the next 10 minutes after induction, suggesting that physical agitation forces are not necessary to induce aggregation **(Fig. 2C, Fig. S1C, Movie S2)**. In fact, *Capsaspora* aggregates are often randomly observed in very dense stationary cultures (35). Agitation may encourage *Capsaspora* aggregation by preventing amoeboid cells from attaching to tissue culture-treated surfaces, therefore leading to greater opportunities for cell-cell contact, adhesion, and ultimately aggregation, as observed in sponges (45, 46). To test this hypothesis, we analyzed the effect of FBS in a dose-response experiment using “ultra-low attachment” surfaces in the absence of agitation **(Fig. 2D, Fig. S1D, Fig. S3, Movie S3, Movie S4)**. Indeed, we found that *Capsaspora* cells aggregate at even 1% (v/v) FBS in the absence of agitation, compared to ∼3% (v/v) FBS in agitation conditions, lending support to the hypothesis that aggregation is primarily regulated by the presence of chemical cue(s) in FBS **(Fig. 2D-F, Fig. S1B and S1D, Movie S4)**. In the low-attachment plates, cells aggregated 10 minutes after induction **(Fig. 2E-F, Fig. S3, Movie S3, Movie S4)**. Upon induction, single cells aggregated into small round aggregates **(Fig. 2E-F, Fig. S3, Movie S3, Movie S4)**. In the next few hours, small aggregates began fusing together into larger aggregates (agglomeration), a process that lasted ∼24 hours **(Fig. 2E-F, Fig. S3, Movie S3)**. After ∼26 hours, all aggregates disassembled into single cells again (disaggregation) **(Fig. 2E, Fig. S3, Movie S3, Movie S4)**. Altogether, these data confirmed that agitation is not necessary to induce aggregation, but rather helps make the cells more sensitive to the chemical inducer(s) in FBS, which are sufficient to induce aggregation in *Capsaspora*.

### Chemical induction of aggregation is reversible

After finding that chemical factor(s) in FBS are necessary to induce aggregation in *Capsaspora*, we asked whether FBS can induce re-aggregation of single cells coming from disassembled aggregates. In fact, re-aggregation of independent cells after tissue dissociation is a common phenomenon frequently observed in sponge cells, hydra cells, and other animal cell lines which, in some cases, is also promoted by serum components (45–47). To this end, we first monitored the aggregation-disaggregation process of *Capsaspora* cells induced with 10% (v/v) FBS **(Fig. 2G, Fig. S4, Movie S5, Fig. S1B-C)**. Then, once the aggregates were completely disaggregated into single-cells, we examined the effect of re-addition of 10% or 30% (v/v) of FBS. Indeed, we found that chemical induction rapidly stimulated re-aggregation of cells within 10 minutes, confirming that *Capsaspora* aggregation is reversible **(Fig. 2G, Fig. S4, Movie S5)**.

### Two soluble FBS components (one small, one large) are required for aggregation

To identify the serum component(s) that induce *Capsaspora* aggregation and re-aggregation, we performed an activity-guided fractionation of FBS. We first used a 30 kDa MW cutoff spin filter to separate FBS into small (<30 kDa) and large (>30 kDa) serum components. These two fractions were then tested for their ability to induce aggregation using the aggregation assay with orbital agitation **(Fig. 2H, Fig. S1B)**. Surprisingly, neither serum fraction was sufficient to induce aggregation alone. However, when both fractions were recombined, the combination induced *Capsaspora* aggregation. These results indicated that at least two soluble serum components are required for aggregation, one that is <30 kDa, and one that is >30 kDa **(Fig. 2H)**.

In parallel, we also assessed the re-aggregation activity of both FBS fractions after disaggregation **(Fig. 2I, Fig. S1B)**. Interestingly, we found that the >30 kDa FBS fraction alone was sufficient to induce re-aggregation. This result suggested that the large FBS fraction component(s) were depleted over time, leading to disaggregation **(Fig. 2I, Fig. S1B)**. Then, re-addition of the >30 kDa component(s) triggered re-aggregation. In contrast, the <30 kDa component(s) likely persisted through the aggregation-disaggregation cycle.

### The “small” active serum component is calcium ions

We then sought to determine the chemical nature of the small active serum component(s). We further used a 3 kDa MW cutoff filter to isolate even smaller serum components, and we found that this <3 kDa FBS fraction induced *Capsaspora* aggregation in the presence of the >30 kDa FBS material (not shown). Next, we performed solvent extractions from freeze-dried <3 kDa FBS material. The solvents included a series of chloroform, methanol and finally water, in order to separate organic-soluble components (such as lipids and polar metabolites) and water-soluble and -insoluble salts. After testing the aggregation activity of each of the recovered extracts in combination with the >30 kDa FBS material, we found that the only active fraction was the one containing salts that did not re-dissolve in any solvent (chloroform, methanol, or water) **(Fig. S5)**.

We then asked which factor(s) in the insoluble fraction could be active. Calcium ions are prevalent in FBS (∼3 mM) (48), and they have the potential to form insoluble salts. Indeed, our FBS contained 3.7 ± 0.2 mM Ca^2+^. We thus hypothesized that calcium ions could be the necessary component for aggregation because of their roles in promoting cadherin-mediated cell-cell adhesion in many organisms (49) and promoting cellular aggregation in sponges, yeasts, and other unicellular species (50, 42, 51). We therefore tested the ability of CaCl2 to induce *Capsaspora* aggregation in the presence of the >30 kDa FBS fraction in a dose-response experiment. Indeed, Ca^2+^ was able to induce *Capsaspora* aggregation at concentrations lower than 0.1 mM **(Fig. 3A).** Therefore Ca^2+^ is sufficient to induce aggregation, when combined with >30 kDa FBS components.

**Figure 3.**
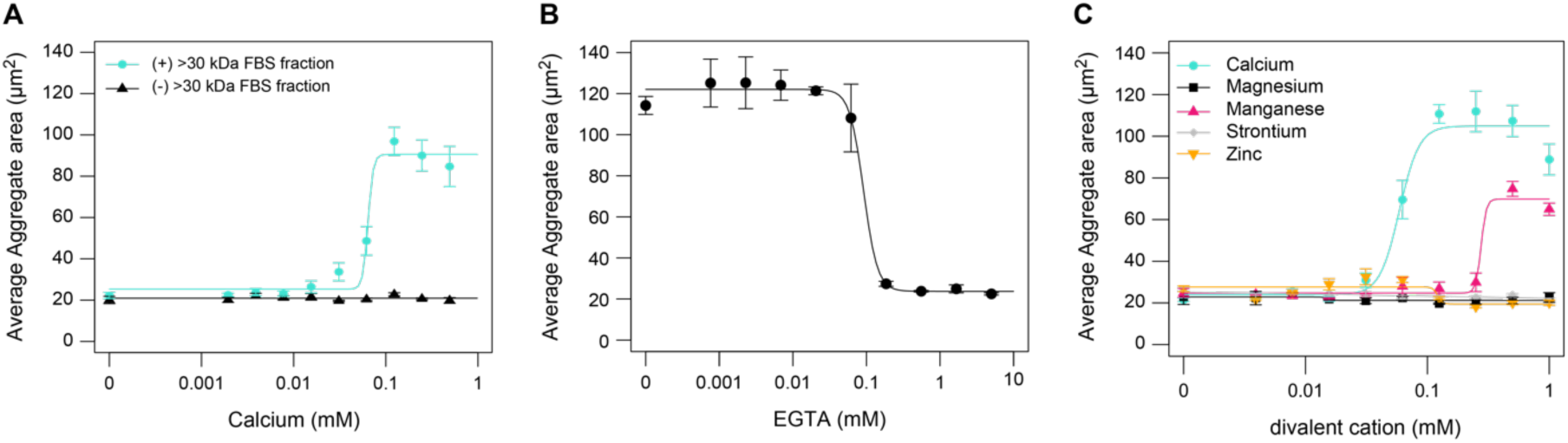
The small (<30 kDa) serum active component is calcium ions. (A) Calcium ions in combination with 5% (v/v) >30 kDa FBS fraction induce aggregation in *Capsaspora*. (B) Treatment with calcium-chelating agent EGTA impairs aggregation of *Capsaspora* cells induced with 5% (v/v) FBS. (C) *Capsaspora* cells induced with 5% (v/v) >30 kDa FBS fraction and chloride salts of various divalent cations aggregate only in the presence of calcium chloride at physiological concentration ranges. Manganese chloride induces weaker aggregation in the presence of 5% (v/v) >30 kDa FBS fraction at high concentrations (0.3 mM). Results in A-C are shown as the average aggregate area of three independent experiments ± s.e.m. All experiments are performed following **Fig. S1D assay**. Figure related to **Fig. S5**.

To test if Ca^2+^ is strictly necessary for aggregation in *Capsaspora*, we depleted available Ca^2+^ with EGTA – a chelating agent with especially high affinity for calcium ions. We performed an EGTA dose-response experiment with *Capsaspora* cells induced with FBS and found that EGTA impaired aggregation at concentrations as low as ∼0.1 mM **(Fig. 3B)**. To rule out the involvement of other divalent metal cations, we tested the activity of the chloride salts of magnesium, manganese, strontium and zinc. None of these cations were capable of inducing aggregation at physiological concentrations **(Fig. 3C).** We observed weak aggregation induced by MnCl_2_, but only at concentrations of 0.3 mM and above, which is far higher than 10% (v/v) FBS would afford (∼0.1 µM, see (48, 52)) **(Fig. 3C)**. Altogether, these findings confirmed that calcium ions are necessary and sufficient to trigger aggregation of *Capsaspora* cells in the presence of >30 kDa FBS components.

### The combination of low-density lipoproteins (LDLs) and calcium ions is sufficient to induce aggregation

Next, we sought to uncover the chemical nature of the large active serum component(s). The aggregation activity of the >30 kDa FBS fraction was sensitive to protease treatment, suggesting that the large active serum component(s) is proteinaceous **(Fig. 4A)**. We therefore separated all proteins from the >30 kDa FBS material by ammonium sulfate precipitation and tested the aggregation activity of the recovered precipitates in combination with 0.2 mM CaCl_2_ **(Fig. S6)**. The resulting active fractions from the ammonium sulfate precipitation **(Fig. S6D-E)** were next separated by anion exchange chromatography and tested for their aggregation activity again **(Fig. S7)**. The active anion exchange fractions **(Fig. S7B)** were in turn further separated by size exclusion chromatography **(Fig. S8)**. The active size exclusion fractions that exhibited robust aggregation **(Fig. S8C)** were later separated by SDS-PAGE and analyzed by Coomassie staining **(Fig. S9A-B)**. We finally determined the protein content of all visible unique bands from the active fractions via mass spectrometry. The major protein in all 7 purified bands matched a single ∼515 kDa protein – Apolipoprotein B100 (ApoB100) **(Fig. S9A-B, Table S1)**.

**Figure 4.**
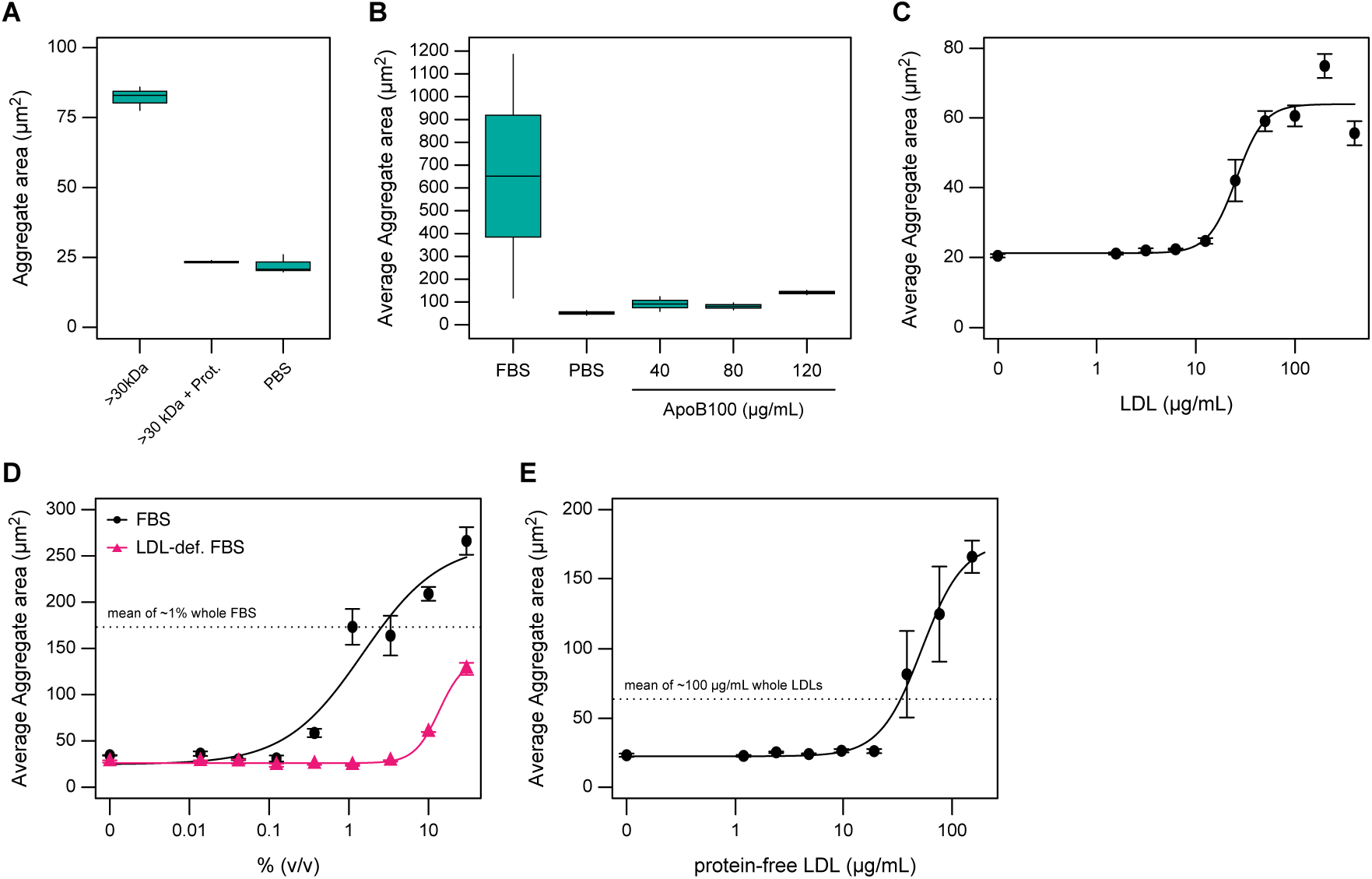
The combination of LDLs and calcium ions induces aggregation in *Capsaspora*. (A) 5% (v/v) of proteinase-treated >30 kDa FBS fraction shows reduced aggregation activity compared to 5% (v/v) of untreated >30 kDa fraction in combination with 0.2 mM CaCl_2_. A 10% (v/v) treatment of 1X PBS was used as negative control. Data are represented as boxplots, showing the median aggregate area (thick black bar) and interquartile ranges from three independent biological replicates. Figure related to **Fig. S6-S8**. (B) ApoB100 in combination with 0.3 mM CaCl_2_ does not fully recapitulate the aggregation activity of 10% (v/v) FBS. Data are represented as boxplots, showing the median aggregate area (thick black bar) and interquartile ranges from two independent biological replicates in agitation conditions **(Fig. S1B assay)**. A 10% (v/v) of FBS was used as a positive control and a 10% (v/v) of 1X PBS was used as a negative control. Figure related to **Fig. S9** and **Table S1**. (C) Dose-response of whole LDLs (plus 0.2 mM CaCl_2_) in the absence of agitation conditions **(Fig. S1D assay)**. Data are represented as the average aggregate area ± s.e.m. from three independent biological replicates. (D) Dose-responses of whole FBS and LDL-deficient FBS (both in the presence of 0.2 mM CaCl_2_) in the absence of agitation **(Fig. S1D assay)**. LDL-deficient FBS has reduced activity, inducing weak aggregation at 30% (v/v) concentrations, whereas whole FBS induces aggregation at ∼1% (v/v). Data are represented as the average aggregate area ± s.e.m. from three independent biological replicates. Figure related to **Fig. S10.** (E) Dose-response of protein-free LDL particles (plus 0.2 mM CaCl_2_) in the absence of agitation conditions **(Fig. S1D assay)**. Data are represented as the average aggregate area ± s.e.m. from three independent biological replicates.

ApoB100 is the primary organizing protein of low-density lipoproteins (LDLs) and very low-density lipoproteins (VLDLs), two major transporters of poorly soluble lipids, including cholesterol, cholesteryl esters, and triglycerides (53, 54). We therefore hypothesized that either ApoB100, itself, induces aggregation or whole lipoprotein particles induce aggregation. Pure ApoB100 (plus 0.3 mM CaCl_2_) failed to induce robust aggregation **(Fig. 4B, Fig. S9C)**. In fact, ApoB100 appeared toxic at high concentrations **(Fig. S9C)**, in agreement with previous work using mammalian cells (55). However, whole LDLs (plus 0.2 mM CaCl_2_) were sufficient to induce aggregation at concentrations as low as ∼10 µg/mL, recapitulating the aggregation activity of our 10% (v/v) FBS control **(Fig. 4C)**. To further confirm the aggregation activity of LDLs, we obtained serum that was >95% deficient in lipoproteins, including LDLs **(Fig. S10A-C)**. We found that LDL-deficient FBS was a substantially less potent aggregation inducer than whole FBS **(Fig. 4D, Fig. S10D)**, confirming the importance of LDLs to induce aggregation.

### Lipids are the active components of low-density lipoproteins

Our finding that *Capsaspora* aggregation is induced by whole LDLs left two major questions. First, LDLs are complexes isolated from natural sources, which contain small amounts of impurities. Therefore, the question remained whether the induction response was truly due to LDLs or instead caused by slight impurities that associate with LDLs. Second, the ecological relevance of LDLs for *Capsaspora* aggregation remained cryptic, especially because LDLs are a vertebrate innovation (56) that *Capsaspora* might not encounter in its natural environment. In fact, *Capsaspora* has only been isolated from snails (57, 58), and its common ancestors with animals would have existed in a world without LDLs. Thus, LDLs are unlikely to be a relevant aggregation cue for present-day *Capsaspora* and its pre-metazoan ancestors. Therefore, the question remained if a more ecologically relevant aggregation inducer exists. To address both of these questions, we sought to determine if pure and ubiquitous components of LDLs could induce aggregation.

Given that the pure ApoB protein failed to induce *Capsaspora* aggregates like whole LDLs, we hypothesized that the lipid components of LDLs could be responsible for inducing aggregation (in combination with calcium ions). We generated protein-free LDL particles (pfLDLs) composed of synthetic lipids representing the 5 major lipid classes in LDLs (50% 1-palmitoyl-2-oleoyl-glycero-3-phosphocholine [POPC]; 5% oleoyl sphingomyelin; 13% cholesterol; 4% triolein; 28% cholesteryl oleate) (59). Transmission electron microscopy images and dynamic light scattering measurements confirmed the formation of lipid particles with an average diameter of 105 nm **(Fig. S11)**, and LC-MS/MS analysis confirmed the incorporation of all the lipids at the percentages listed above **(Table S2)**. Indeed, these pfLDLs were sufficient to induce aggregation (in the presence of calcium ions) **(Fig. 4E)**. The EC_50_ of induction was 53 µg/mL, which is comparable to the potency of LDLs (EC_50_ ∼26 µg/mL). Therefore, one or more of the polar lipids of LDLs appear to be the true aggregation cue.

### Dynamic lipoprotein concentration regulates the aggregative state of *Capsaspora*

Given that aggregates dissociated over time (and re-formed after addition of the lipoprotein-containing serum fractions), we inquired whether LDLs were depleted over the course of the aggregation experiment. To test this, we analyzed the LDL content during the course of aggregation and disaggregation using an anti-LDL antibody. Our findings revealed that the concentration of LDL did, in fact, significantly decrease in the time leading up to disaggregation **(Fig. 5A, Fig. S12)**. These results led us to hypothesize that the LDLs were depleted by *Capsaspora* metabolism. To test this hypothesis, we assessed the localization of LDLs using immunofluorescence microscopy. We observed LDLs localized in intracellular vesicles in *Capsaspora* aggregates and adherent cells immunostained with anti-LDL antibodies **(Fig. 5B, Fig. S13).** Together, these findings indicate that *Capsaspora* engulfs LDLs, thereby depleting them from the media. Therefore, the dynamic concentration of lipoproteins regulates the aggregative state of *Capsaspora*: high concentrations of lipoproteins induce aggregation; however, as lipoproteins are depleted, aggregates disassemble and only re-assemble when lipoprotein concentrations are restored.

**Figure 5.**
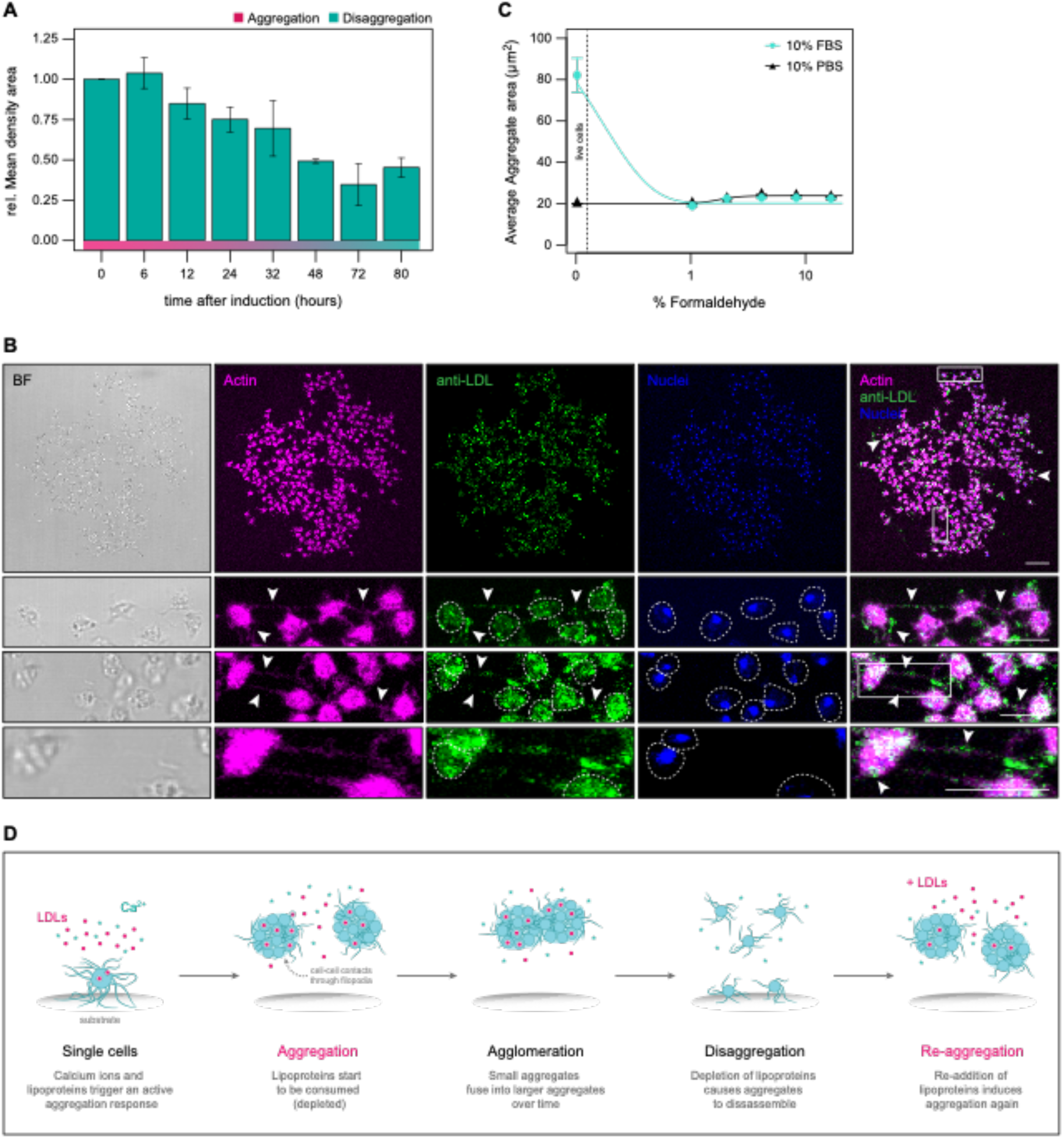
Dynamic lipoprotein concentration regulates an aggregation response in viable *Capsaspora* cells. (A) LDLs are depleted over time during the aggregation-disaggregation process. Quantification of LDL content of media supernatants collected from cells induced with 10% (v/v) FBS (0h) by Western Blot show progressive disappearance of LDLs in the time leading up to disaggregation (72-80h). Data are represented as the mean density areas relative to 0h timepoint ± s.d. from two independent biological replicates. Figure related to **Fig. S4A-B** and **Fig. S12**. (B) LDLs are localized in intracellular small foci and in patches along filopodia in a *Capsaspora* aggregate induced with 10% (v/v) of FBS. Shown is a maximum z-projection of an aggregate (upper panel) stained with phalloidin (magenta) to mark filamentous actin (cell body and filopodia), an anti-LDL antibody to mark LDLs (green), and DRAQ5 to mark nuclei (blue). A merge of phalloidin and anti-LDL staining shows that LDLs localize in the interior of cells and filopodia (white arrows). A representative brightfield (BF) stack image is shown. Lower panels represent two zoom-in maximum z-projection images (white squares) of the aggregate stained as before. Scale bar in the upper panel represents 25 µm and 10 µm in all magnified lower panels. Figure related to **Fig. S13.** (C) *Capsaspora* cells fixed with increasing concentrations of formaldehyde do not aggregate upon induction with 10% (v/v) of FBS. Data are represented as the average aggregate area ± s.e.m. from three independent biological replicates **(Fig. S1B assay)**. A 10% (v/v) of 1X PBS was used as negative control. (D) *Capsaspora* cells aggregate in the presence of calcium ions and lipoproteins, which trigger an active aggregation response. During the course of aggregation, smaller aggregates fuse together into larger aggregates (agglomeration) while consuming lipoproteins. Depletion of lipoproteins leads aggregates to disassemble into single cells again (disaggregation). Re-addition of lipoproteins induces re-aggregation of *Capsaspora* cells.

### Only viable *Capsaspora* cells aggregate

Finally, we began to dissect the mechanism by which the active components from FBS induce aggregation in *Capsaspora*. In sponges, cellular aggregation is induced by soluble proteoglycan complexes called sponge *aggregation factors* (AFs) (43, 44, 50, 60). These aggregation inducers function as multivalent adhesives to physically link sponge cells to each other, even when those cells are formalin-fixed (*i.e.,* metabolically inactive) in rotationally-induced aggregation experiments (45, 60). In other unicellular species, like *D. discoideum,* yeasts or algae, the cues induce an active cellular response that results in aggregation (40–42). Actin staining revealed that cells within *Capsaspora* aggregates appear directly linked to each other through filopodia **(Fig. 5B, Fig. S13)**. Because we also observed distinct patches of LDLs localized along filopodia within aggregates **(Fig. 5B, Fig. S13),** we inquired whether serum-supplied lipoproteins trigger *Capsaspora* aggregation by an adhesive mechanism like sponge AFs. We thus fixed *Capsaspora* cells with increasing concentrations of formaldehyde and later induced aggregation with 10% (v/v) FBS in orbital agitation **(Fig. 5C, Fig. S1B)**. Unlike sponge cells, we found that formalinized *Capsaspora* cells were *unable* to aggregate even in the presence of 10% (v/v) FBS. Although we cannot rule out the possibility that formaldehyde treatment directly disrupts an LDL-*Capsaspora* adhesive interaction, this result suggests that LDLs are *not simply a “glue”* that links passive cells together. Instead, LDLs likely induce aggregation through an active biological response from living *Capsaspora* cells, in a manner reminiscent to the active cellular responses observed in other unicellular species.

## DISCUSSION

We have discovered that chemical cues induce aggregative multicellularity in *Capsaspora owczarzaki,* one of the closest unicellular relatives of animals **(Fig. 5D)**. In the presence of these cues, *Capsaspora* amoebae form large spherical aggregates that resemble the initial aggregated stages whereby dissociated animal cells re-aggregate and develop into new viable organisms (17, 23–25, 27, 29). Thus, the fact that early-branching animals, the choanoflagellate *S. rosetta* (39) and *Capsaspora* utilize chemical cues to regulate aggregation is suggestive that the unicellular ancestors of animals also employed chemical cues to regulate aggregation.

The first chemical factors that we found necessary for *Capsaspora* aggregation are calcium ions (corroborated by recent work from Phillips *et al.,* (61)). Calcium ions are similarly required for the aggregation of *D. discoideum,* yeasts and sponge cells. During *D. discoideum* aggregation, intracellular calcium ion dynamics are linked to cAMP levels, that in turn regulate chemotaxis and stalk cell differentiation (51). Furthermore, calcium ions act as secondary messengers in many signaling pathways (62) and may be required for transduction of the *Capsaspora* aggregation-induction signal. Alternatively, calcium ions are essential for stabilizing many cell-cell adhesion molecules. For example, calcium ions stabilize the lectins responsible for yeast asexual flocculation (42). In sponges, calcium ions likely play a structural role stabilizing cell-cell contacts through associations of cell-surface sponge *aggregation factors* (AFs) (43, 44, 50, 60). Calcium ions also stabilize other animal cell adhesion molecules, such as cadherins and integrins (63, 64). Indeed, some genes predicted to encode calcium-dependent adhesion proteins are upregulated in *Capsaspora* aggregates (35, 37, 38). Therefore, the requirement of Ca^2+^ for *Capsaspora* aggregation may be to stabilize these adhesion molecules. In fact, we observed several cell-cell contacts between filopodia of neighboring cells within aggregates, which may be the loci of these calcium-dependent cell-cell adhesion complexes. The exact mechanism of calcium-dependent aggregation is a topic of further research, but its general requirement is a similarity of aggregation induction in *Capsaspora* and sponges, as well as other animals and unicellular species.

The second chemical factor necessary to induce *Capsaspora* aggregation is a low-density lipoprotein (LDL), particularly the lipids present in LDLs. This inducer has similarities and differences with the aggregation inducers of other organisms – notably in their chemical composition, their source (environmental vs. organism-produced), and their general mechanism of action (signal vs. adhesive). Regarding the *composition* of the inducer itself, we are not aware of any organisms whose aggregation is induced by whole LDLs. LDLs are a similar size to sponge AFs, which are large soluble proteoglycans (43, 44, 60). This size affords the potential for multivalent interactions with the surfaces of one or more cells. In addition, the fact that lipids present in LDLs are the *Capsaspora* aggregation inducers also demonstrates some similarity with lipid kairomones that induce aggregation in some algae (40). In contrast, the inducer of choanoflagellate mating aggregation is a large protein enzyme (39), and *Dictyostelium* and yeast aggregation are regulated by small molecules and peptides (41, 42). Regarding the *source* of the inducer, we have seen no evidence that *Capsaspora* secretes its aggregation inducer. Instead, it is likely an environmental cue, as are the inducers of aggregation in the choanoflagellate *S. rosetta* and some algae (39, 40). Likewise, the *absence* of environmental nutrients plays a role in triggering aggregation in yeast and *Dictyostelium*. In contrast, sponge AFs, as well as yeast pheromones and *Dictyostelium* cAMP are signals produced by the organism, itself. Finally, in terms of *mechanism*, it appears that only viable *Capsaspora* cells are capable of aggregating in response to lipoproteins and calcium ions. This finding suggests that these cues trigger an active cellular response, like cues do in yeast, *Dictyostelium* and algae – unlike sponge AFs, which act as adhesives to link even formaldehyde-fixed cells. Therefore, although chemically regulated aggregation occurs in some animals, unicellular holozoans, and other unicellular eukaryotes, there is a wide range of inducers, sources, and mechanisms responsible.

The ecological and evolutionary relevance of lipoprotein-induced aggregation in *Capsaspora* is under additional investigation. *Capsaspora* has, so far, only been found inside snails (57, 58, 65), which do not possess classical LDLs or other types of vertebrate lipoproteins (56, 66). Therefore, it is possible that the LDL-mediated aggregation response is not relevant in *Capsaspora*’s natural environment. However, we found that reconstituted particles of sphingomyelin, phosphatidylcholine, cholesterol, triglyceride, and cholesteryl ester induced aggregation in *Capsaspora*. These lipids are common in animals, including *Biomphalaria* snails (67, 68). Therefore, it is plausible that *Capsaspora* aggregation is an evolved response to chemical cues from the snail host environment. Alternatively, *Capsaspora* may also live outside snail hosts where extracellular vesicles or other lipid/protein complexes from neighboring microbes may induce an aggregative behavior. In this case, the lipoprotein response could be ancient, even predating animals, as lipid signals are also known to trigger multicellular responses in other unicellular species (7, 40, 69). Indeed, phosphatidylcholines and sterols are commonly produced by microbial eukaryotes (70, 71), and some bacteria produce phosphatidylcholines and triglycerides as well (72, 73). Additionally, aggregation may be induced by other polar lipids that are not prevalent in LDLs but are ubiquitous in bacterial membranes (e.g., phosphatidylglycerols, phosphatidylethanolamines, cardiolipins). Therefore, in ongoing work, we are comprehensively investigating which lipid classes induce aggregation, and whether these components are also present in snail hosts or released by environmental microbes.

Taken together, the prevalence of chemically regulated aggregation, particularly in unicellular holozoans and some early branching animals suggests that it is a broadly beneficial phenotype and therefore plausibly existed in the recent unicellular ancestors of animals. A remaining question is which mechanisms likely underpinned these phenotypes in their common unicellular ancestors – and if they continued to play important functions in the first animals. A route to address these questions is to characterize the genetic basis by which aggregation is regulated in *Capsaspora* and other unicellular holozoans and compare these to pathways employed in extant animals. If homologous genes are employed, then those steps could imply ancient conserved mechanisms of a regulated aggregation pathway. Importantly, our work identifying the inducers of aggregation will enable these essential investigations of the signaling pathways that initiate the *Capsaspora* aggregation process.

## CONCLUSION

We discovered that the aggregative behavior of *Capsaspora* is a reversible process induced by a combination of calcium ions and lipids present in lipoproteins. Therefore, extant animals, other unicellular species, and this close unicellular relative to animals utilize chemical cues to regulate reversible aggregation. Similar to sponges, other animal species, and more distantly-related unicellular organisms (*e.g.,* yeasts and *D. discoideum*)*, Capsaspora* requires calcium ions for aggregation. Although lipoproteins seem to be *Capsaspora*-specific cues, they resemble sponge AFs in their size and potential for multivalent interactions. Further characterization of the aggregation induction pathway in *Capsaspora* is warranted to determine the degree of conservation with extant animals. The results will help determine the likelihood that a regulated aggregation mechanism was a trait of the first animals. Furthermore, our findings combine with other recent studies to reveal a trend that unicellular holozoans utilize chemical triggers to regulate diverse multicellular phenotypes (39, 69). Although more taxa must be examined, these observations hint that soluble chemical cues were important in regulating reversible multicellularity in the close ancestors of animals.

## MATERIALS AND METHODS

### Cell strain and growth conditions

*Capsaspora owczarzaki* cell cultures (strain ATCC®30864) were grown axenically in 25 cm^2^ culture flasks (#734-0044, Falcon® VWR) with 5 mL ATCC media 1034 (modified PYNFH medium) containing 10% (v/v) heat-inactivated Fetal Bovine Serum (FBS) (#F9665-100mL, Sigma-Aldrich), hereafter *growth media*, in a 23°C incubator. Adherent stage cells (filopodiated amoebae) at the exponential growth phase were obtained by passaging ∼100-150 µL of adherent cells at ∼90% confluence in 5 mL of growth media and grown for 24-48h at 23°C.

### Fetal Bovine Serum contains chemical factor(s) necessary for aggregation in *Capsaspora*

#### Media lacking components experiment (related to Fig. 2A-B and Fig. S1A)

Adherent cells from a ∼90% confluence culture were scraped, homogenized and counted to seed 7.5*10^5^ cells in 1 mL of growth media per well in a 12-well plate (#55428, Nunc/DDBioLab) and incubated overnight at 23°C. After incubation, each well was washed once with 1 mL of either peptone-free media; yeast extract-free media; FBS-free media, 1X PBS or growth media, and later filled with 1 mL of the corresponding media. Next, cells were put at 50 rpm orbital agitation (Celltron Bench-Top Shaker, INFORS-HT) during 24h at room temperature. Aggregates were imaged after 24h agitation at 36 distinct locations throughout each well at 5X magnification using transmitted light (brightfield) on a Zeiss Axio Observer Z.1 epifluorescence inverted microscope equipped with LED illumination and an Axiocam 503 mono camera. All experiments were performed in three biological replicates, testing each condition per duplicate.

### Quantification of average aggregate area (related to Fig. S2)

Images were batch processed in Fiji Imaging Software version 2.1.0/1.53c (74) using a macro script. In brief, images were first converted to binary using the ‘Make Binary’ command in the “Process” menu using the default method. Images were then analyzed using the ‘Analyze Particles’ command to calculate the area of each particle, setting the analysis parameters to a particle size range of 10-infinity µm^2^ and a circularity of 0.01-1, excluding particles on the edges of the images. A minimum of 2000 particle areas are quantified in each condition.

An extra “Subtract Background” step was added to reduce shadows in images from **Fig. 2B** and **Movies S1, S2, S4** and **S5**. The parameters were set to a rolling ball radius to 20 pixels **(Fig. 2B** and **Movies S1, S2** and **S5)** or 100 pixels **(Movie S4)** and selecting the sliding paraboloid option in all of them. After this step, images were batch processed and analyzed using a macro script as before.

### Chemical induction is sufficient for aggregation, even in the absence of physical agitation

#### Aggregation dynamics over time in the absence of orbital agitation (related to Fig. 2C, Fig. S1C, Movie S1 and Movie S2)

Adherent stage cells from a ∼90% confluence culture were scraped, homogenized and harvested at 5000xg during 5 min at room temperature. Cells were washed twice with 10 mL of FBS-free media and later resuspended to an appropriate volume of FBS-free media to reach a cell concentration of 8.33*10^6^ cells/mL. Next, 7.5*10^5^ cells were seeded in either 900 µL (10% (v/v) FBS condition) or 700 µL (30% (v/v) FBS condition) of FBS-free media per well in a 12-well plate (#55428, Nunc/DDBioLab) and starved at least 14h at 23°C. After starvation, cells were imaged at 4 distinct locations throughout each well every 3 min for 22 hours at 5X magnification using transmitted light (brightfield) and a 610 nm LongPass Color filter (#FGL610, Thorlabs) on a Zeiss Axio Observer Z.1 epifluorescence inverted microscope equipped with LED illumination and an Axiocam 503 mono camera. Aggregation was induced by the addition of either a 10% (v/v) or a 30% (v/v) of FBS added at the top wall of each well after 9 min of imaging. The corresponding volume of 1X PBS was used as negative control. All experiments were performed in three biological replicates, testing each condition per duplicate.

#### FBS-dose response experiment in orbital agitation (related to Fig. 2D and Fig. S1B)

Adherent stage cells from a ∼90% confluence culture were scraped, homogenized and harvested at 5000xg during 5 min at room temperature. Cells were washed twice with 10 mL of FBS-free media and later resuspended to an appropriate volume of FBS-free media to reach a cell concentration of 8.33*10^6^ cells/mL. Next, 7.5*10^5^ cells were seeded in 900 µL of FBS-free media per well in a 12-well plate (#55428, Nunc/DDBioLab) and starved at least 14h at 23°C. After starvation, 0-30% (v/v) of FBS was added in each well in a 1:3 serial dilution. Cells were immediately put at 50 rpm orbital agitation at room temperature (Celltron Bench-Top Shaker, INFORS-HT). Aggregates were imaged after 24h agitation at 10 distinct locations throughout each well at 10X magnification using transmitted light (brightfield) on an Eclipse TS100 Nikon epifluorescence inverted microscope equipped with an Intensilight C-HGFI Illuminator and a DS-Fi2 Camera Head. All experiments were performed in three biological replicates, testing each condition per duplicate.

#### FBS-dose response experiment in the absence of orbital agitation (related to Fig. 2D-F, Fig. S1D, Fig. S3, Movie S3 and Movie S4)

Adherent stage cells from a 2-day grown ∼90% confluence culture were scraped, homogenized and washed once with 15 mL of FBS-free media and allowed to starve overnight in a tube at 23°C. After starvation, 8*10^5^ cells were seeded in 150 µL of FBS-free media per well in a 96-well ultra-low attachment microplate (#CLS3474, Corning) and allowed to settle for 2 hours at room temperature. To assess the effect of FBS in a dose response experiment, 50 µL containing 0-30% (v/v) of FBS was added in each well in a 1:3 serial dilution (in FBS-free media). Aggregates were imaged 90 min after induction at 5X magnification using transmitted light (brightfield) in a Leica DMi1 inverted microscope **(Fig. 2D)**. To monitor the dynamics of aggregation over time, 50 µL of either 0%, 0.5%, 1.5%, 5% or 10% (v/v) of FBS were added in each well and imaged every 10 minutes for 42h at 5X magnification using transmitted light (brightfield) in an A1 Nikon inverted microscope with an automatic stage **(Fig. 2E-F, Fig. S3, Movie S3** and **Movie S4).** All experiments were performed in three biological replicates.

### Chemical induction of aggregation is reversible

#### Re-aggregation experiment in orbital agitation (related to Fig. S1B and Fig. S4)

Adherent cells from a ∼90% confluence culture were scraped, homogenized and harvested at 5000xg during 5 min at room temperature. Cells were washed twice with 10 mL of FBS-free media and later resuspended to an appropriate volume of FBS-free media to reach a cell concentration of 8.33*10^6^ cells/mL. Next, 7.5*10^5^ cells were seeded in 900 µL of FBS-free media per well in a 12-well plate (#55428, Nunc/DDBioLab) and starved at least 14h at 23°C. After starvation, 10% (v/v) of FBS was added per well and cells were immediately put in 50 rpm orbital agitation (Celltron Bench-Top Shaker, INFORS-HT) at room temperature. Aggregation and disaggregation of cells was periodically monitored during 80h. Once 100% of cells were disaggregated (after ∼80h agitation) re-aggregation was induced by re-adding 10% (v/v) FBS per well. Samples supernatant from all timepoints were collected to be further analyzed by Western Blot (see below). Aggregates were imaged at 10 distinct locations throughout each well at 10X magnification using transmitted light (brightfield) on an Eclipse TS100 Nikon epifluorescence inverted microscope equipped with an Intensilight C-HGFI Illuminator and a DS-Fi2 Camera Head. All experiments were performed in three biological replicates, testing each condition per duplicate.

#### Re-aggregation experiment in the absence of orbital agitation (related to Fig. 2G, Fig. S1C and Movie S5)

Adherent stage cells from a ∼90% confluence culture were scraped, homogenized and harvested at 5000xg during 5 min at room temperature. Cells were washed twice with 10 mL of FBS-free media and later resuspended to an appropriate volume of FBS-free media to reach a cell concentration of 8.33*10^6^ cells/mL. Next, 7.5*10^5^ cells were seeded in 900 µL of FBS-free media per well in a 12-well plate (#55428, Nunc/DDBioLab) and starved at least 14h at 23°C. After starvation, cells were imaged at 8 distinct locations throughout each well every 3 min for 26 hours at 5X magnification using transmitted light (brightfield) and a 610 nm LongPass Color filter (#FGL610, Thorlabs) on a Zeiss Axio Observer Z.1 epifluorescence inverted microscope equipped with LED illumination and an Axiocam 503 mono camera. Aggregation was induced by the addition of 10% (v/v) of FBS added at the top wall of each well after 9 min of imaging. Cells were monitored until 100% of the aggregates were disaggregated into single-cells again (around 19h after induction). At this point, an additional 30% (v/v) FBS or 30% (v/v) of 1X PBS (negative control) were re-added as before. All experiments were performed in three biological replicates.

### Two soluble FBS components (one small, one large) are required for aggregation

#### Effect of FBS fractions on aggregation (related to Fig. 2H and Fig. S1B)

FBS was separated into >30 kDa and <30 kDa fractions with an Amicon Ultra 30 kDa MW cutoff centrifugal filter (#UFC903024, Millipore). The <30 kDa flow-through was retained for testing, and the >30 kDa material was washed three times with PBS to minimize residual <30 kDa FBS material. In parallel, adherent stage cells from a ∼90% confluence culture were scraped, homogenized and harvested at 5000xg during 5 min at room temperature. Cells were washed twice with 10 mL of FBS-free media and later resuspended to an appropriate volume of FBS-free media to reach a cell concentration of 2*10^7^ cells/mL and allowed to starve overnight at 23°C. Next, 4*10^6^ cells were seeded in 2 mL of FBS-free media per well in a 12-well plate (#55428, Nunc/DDBioLab) and a 10% (v/v) of inducers (whole FBS, <30 kDa or >30 kDa FBS fractions, a combination of a 5% (v/v) of <30 kDa plus 5% (v/v) of >30 kDa FBS fractions, or 1X PBS control) were added in each well. Cells were immediately put at 50 rpm orbital agitation at room temperature (Celltron Bench-Top Shaker, INFORS-HT). Aggregates were imaged after 24h agitation at 9 distinct locations throughout the well at 5X magnification using transmitted light (brightfield) on a Leica DMi1 inverted microscope. All experiments were performed in three biological replicates.

#### Effect of FBS fractions on re-aggregation (related to Fig. 2I and Fig. S1B)

Adherent stage cells from a ∼90% confluence culture were scraped, homogenized and harvested at 5000xg during 5 min at room temperature. Cells were washed twice with 10 mL of FBS-free media and later resuspended to an appropriate volume of FBS-free media to reach a cell concentration of 2*10^7^ cells/mL and allowed to starve overnight at 23°C. Next, 2*10^6^ cells were seeded in 2 mL of FBS-free media per well in a 12-well plate (#55428, Nunc/DDBioLab) and 10% (v/v) of whole FBS was added in each well. Cells were immediately put at 50 rpm orbital agitation at room temperature (Celltron Bench-Top Shaker, INFORS-HT) until 100% disaggregation (∼72h). At this point, 10% (v/v) of inducers (see above) were added and cells were put back to agitation. Aggregates were imaged after 1-2h agitation upon re-induction at 9 distinct locations throughout the well at 5X magnification using transmitted light (brightfield) on a Leica DMi1 inverted microscope. All experiments were performed in three biological replicates.

### The “small” active serum component is calcium ions

#### Separation of the active <3 kDa FBS components by solubility

Whole FBS (15 mL) was passed through an Amicon Ultra 3 kDa MW cutoff centrifugal filter (#UFC900308, Millipore) and the resulting ∼15 mL that passed through the filter (corresponding to <3 kDa FBS material) was frozen and lyophilized to yield a powder. The resulting powder was added to 1 mL of chloroform (#CX1058-1, EMD Millipore), and the mixture was sonicated and vortexed to dissolve any organic-soluble components. The insoluble precipitate was removed by centrifugation, and the chloroform supernatant was collected and evaporated to dryness. The remaining precipitate was added to 1 mL of 100% methanol (#MX0475-1, EMD Millipore), and the above procedure was repeated to yield a vial of dry methanol-soluble material and remaining insoluble precipitate. This process was repeated serially with 80% methanol diluted in ultrapure MilliQ water, 20% methanol diluted in ultrapure MilliQ water, and finally 100% ultrapure MilliQ water. After this procedure, some precipitate still did not redissolve in the water. The dried supernatants were redissolved as follows: the chloroform supernatant was redissolved in 100 µL of 80/20 chloroform/methanol, the others (100% methanol, 80% methanol, 20% methanol, and water) were redissolved in 1 mL ultrapure MilliQ water and sterile-filtered through 0.22 µM syringe filter (#431224, Corning). The water-insoluble precipitate was resuspended in 1 mL ultrapure MilliQ water.

#### Determination of active <3 kDa FBS fraction (related to Fig. S5)

Each <3 kDa FBS extracted material was added to >30 kDa FBS fraction in 2 mL FBS-free media in the well of a 6-well plate (#10861-554, VWR) at roughly 10X the concentration they would exist in the standard growth media. Specifically, to wells containing confluent adherent cells, 2 mL of FBS-free media was added to each well, followed by 150 µL of >30 kDa FBS components in 1X PBS at their 1X FBS concentrations (so the final concentration of the >30 kDa FBS components in the assay well was comparable to 10% (v/v) FBS being added to the assay). Then, the following volumes of the <3 kDa FBS material were added: chloroform fraction (30 µL), methanol fraction (100 µL), methanol/water fractions (100 µL), water fraction (100 µL), and insoluble water suspension (50 µL). A negative control with only the FBS-free media and >30 kDa FBS components added was also included in the assay. The plates were then gently agitated at 50 rpm orbital shaking at room temperature overnight.

#### Calcium ions quantification in FBS

Calcium ions content in FBS was quantified via a colorimetric assay (#MAK022, Sigma-Aldrich) following manufacturer’s instructions. Results are represented as mean ± s.d. in three replicates.

#### Calcium rescue experiment (related to Fig. 3A and Fig. S1D)

Adherent stage cells from a 2-day grown ∼90% confluence culture were scraped, homogenized and washed once with 15 mL of FBS-free media and allowed to starve overnight in a tube. After starvation, 8*10^5^ cells were seeded in 150 µL of FBS-free media per well in a 96-well ultra-low attachment microplate and allowed to settle for 2 hours at room temperature. In parallel, the >30 kDa FBS fraction was washed 3 times with 1X PBS containing 0.5 M EGTA (#SC-3593B, ChemCruz) adjusted to pH 7 and then washed again 3 times more with 1X PBS to remove excess EGTA. A 5% (v/v) of washed >30 kDa FBS fraction was premixed with a 1:3 serial dilution of CaCl_2_ (#C5080, Sigma-Aldrich) diluted in 1X PBS (concentration range from 0.5 to 0 mM) and later added to each well. A 5% (v/v) of 1X PBS premixed with a 1:3 serial dilution of CaCl_2_ as before as used as negative control. Aggregates were imaged 90 min after induction at 5X magnification using transmitted light (brightfield) in a Leica DMi1 inverted microscope. All experiments were performed in three biological replicates.

#### EGTA inhibition experiment (related to Fig. 3B and Fig. S1D)

Adherent stage cells from a 2-day grown ∼90% confluence culture were scraped, homogenized and washed once with 15 mL of FBS-free media and allowed to starve overnight in a tube at 23°C. After starvation, 8*10^5^ cells were seeded in 150 µL of FBS-free media per well in a 96-well ultra-low attachment microplate and allowed to settle for 2 hours at room temperature. Whole FBS was premixed with a 1:3 serial dilution of EGTA (concentration range from 5 mM to 0 mM) diluted in 1X PBS and adjusted to pH 7 and later added to each well, reaching a final FBS concentration of 5% (v/v). Aggregates were imaged 90 min after induction at 5X magnification using transmitted light (brightfield) in a Leica DMi1 inverted microscope. All experiments were performed in three biological replicates.

#### Divalent cations dose-response experiments (related to Fig. 3C and Fig. S1D)

Adherent stage cells from a 2-day grown ∼90% confluence culture were scraped, homogenized and washed once with 15 mL of FBS-free media and allowed to starve overnight in a tube at 23°C. After starvation, 8*10^5^ cells were seeded in 200 µL of FBS-free media per well in a 96-well ultra-low attachment microplate and allowed to settle for 2 hours at room temperature. A 1:2 serial dilution (concentration range from 1 mM to 0 mM) of Calcium chloride (#C5080, Sigma-Aldrich), Magnesium chloride (#M2393, Sigma-Aldrich), Manganese chloride (#M3634, Sigma-Aldrich), Strontium chloride (#255521, Sigma-Aldrich), and Zinc chloride (#Z0251, Sigma-Aldrich) were added to each well together with 5% (v/v) of >30 kDa FBS fraction, previously prewashed three times in 0.5 M EGTA diluted in 1X PBS and three more times with 1X PBS. Aggregates were imaged 90 min after induction at 5X magnification using transmitted light (brightfield) in a Leica DMi1 inverted microscope. All experiments were performed in three biological replicates.

### The combination of low-density lipoproteins (LDLs) and calcium ions is sufficient to induce aggregation

#### Protease treatment of >30kDa FBS fraction (related to Fig. 4A)

Protease treatment of >30kDa FBS fraction was performed by treating 200 µL of large fraction (previously washed 3 times with 1X PBS) with 10 µL of 20 mg/mL Proteinase-K (#SAE0151, Sigma-Aldrich) at 37°C for 1 hour, with periodic agitation. After incubation, the mix was cooled down at room temperature for 1 hour, inverting periodically. The mix was further treated with 1 mM PMSF (#P7626, Sigma-Aldrich) at room temperature for 4 hours, mixing periodically. The final solution was washed 3 times with 1X PBS using an Amicon Ultra 30 kDa MW cutoff centrifugal filter (#UFC903024, Millipore) to remove unreacted PMSF and then resuspended to 200 µL (same volume as original sample). The aggregation activity of the protease-treated fraction was then tested at 5% (v/v) with 0.2 mM CaCl_2_ supplemented back. A 5% (v/v) of 1X PBS plus 0.2 mM CaCl_2_ were used as a negative control. Aggregates were imaged 90 min after induction at 5X magnification using transmitted light (brightfield) in a Leica DMi1 inverted microscope. All experiments were performed in triplicate.

#### Separation of >30 kDa FBS components by ammonium sulfate solubility (related to Fig. S6)

Whole FBS (500 mL) was concentrated to 80 mL with Amicon Ultra 30 kDa MW cutoff centrifugal filters (#UFC903024, Millipore). This material was split evenly between four centrifuge tubes, each diluted to a final volume of 30 mL and a final concentration of 100 mM Tris-HCl, pH 8.0. To each tube was added 3.18 g ammonium sulfate for a final concentration of 20%. The tubes were rocked in ice for 30 minutes, then centrifuged at 20,000xg for 15 min. The supernatants were transferred to new tubes and the pellets were redissolved in 25 mL of ultrapure MilliQ water. To each supernatant tube was added another 1.65 g of ammonium sulfate to reach a concentration of 30%. The tubes were mixed and centrifuged as before, and the pellets were resuspended as before. This process was repeated for the following concentrations of ammonium sulfate: 40% (+1.68 g), 50% (+1.74 g), 55% (+0.9 g), 60% (+0.9 g), 65% (+0.93 g), 70% (+0.93 g), and 80% (+1.95 g). To test activity of each fraction, 50 µL of each was added to 2 mL cultures of attached cells in 6-well plates (#10861-554, VWR) containing 0.2 mM CaCl_2_. The plates were agitated at 50 rpm at room temperature for 4 hours and aggregation was assessed at 5X magnification using transmitted light (brightfield) in a Leica DMi1 inverted microscope.

#### Separation of active ammonium sulfate precipitate fraction by anion exchange chromatography (related to Fig. S7)

From the material that precipitated at 65% ammonium sulfate, 8 mL was exchanged into 20 mM ethanolamine/acetic acid buffer pH 9.2 and concentrated to 2 mL with an Amicon Ultra 100 kDa MW cutoff centrifugal filter (#UCF5100, Millipore). A HiPrep Q XL 16/10 anion exchanger column (#28936538, Cytiva) was used at a 5 mL/min flow rate to separate this protein mixture using 20 mM ethanolamide/acetic acid buffer pH 9.2 with a gradient from 0 M NaCl (solvent A) to 1 M NaCl (solvent B). The gradient follows: 0-3 min (0% B), 3-38 min (linear gradient to 60% B), 38-40 min (linear gradient to 100% B), 40-45 min (hold 100% B). Fractions were collected every 20 seconds (1.67 mL). Fractions were exchanged into 1X PBS and added (20 µL) to 1 mL cultures of *Capsaspora* cells in wells of 12-well plates in FBS-free media containing 0.25 mM CaCl_2_. Plates were agitated at 50 rpm at room temperature for 1 hour and aggregation was assessed at 5X magnification using transmitted light (brightfield) in a Leica DMi1 inverted microscope.

#### Separation of active anion exchange fractions by size exclusion chromatography (related to Fig. S8)

The most active fractions from the anion exchange column were further separated by size exclusion chromatography. Two adjacent fractions (1 and 2 in **Fig. S8**) were combined, and the third active fraction was kept separate. Both of these fractions were exchanged with 1X PBS pH 6.5 and concentrated to 100 µL with Amicon Ultra 50 kDa MW cutoff centrifugal filters (#UFC5050, Millipore). Each was separated through a Superdex 200 Increase 10/300 GL column (#28990944, Cytiva) with a 0.75 mL/min flow rate in 1X PBS pH 6.5. Fractions were collected every 30 seconds (0.375 mL). Fractions were concentrated to 20 µL with Amicon Ultra 50 kDa MW cutoff centrifugal filters, and 5 µL of each was added to 1 mL cultures of *C. owczarzaki* in wells of 12-well plates in FBS-free media containing 0.25 mM CaCl_2_. Plates were agitated at 50 rpm at room temperature for 1 hour and aggregation was assessed at 5X magnification using transmitted light (brightfield) in a Leica DMi1 inverted microscope.

#### Determination of protein content of active size exclusion fractions via mass spectrometry of tryptic peptides (related to Fig. S9A-B and Table S1)

The most active fractions from the size exclusion column were analyzed by SDS-PAGE and Coomassie staining. All visible unique bands (7 total) were excised and subjected to in-gel trypsin digestion and LC-MS/MS analysis by the Taplin Mass Spectrometry Facility at Harvard Medical School. The major protein in all 7 bands was Apolipoprotein B100 (ApoB100) (see **Table S1**).

#### ApoB100 effect on *Capsaspora* aggregation (related to Fig. 4B, Fig. S1A, Fig. S9C)

Adherent cells from a ∼90% confluence culture were scraped, homogenized and harvested at 5000xg during 5 min at room temperature. Cells were washed twice with 10 mL of FBS-free media and later resuspended to an appropriate volume of FBS-free media to reach a cell concentration of 8.33*10^6^ cells/mL. Next, 7.5*10^5^ cells were seeded in 900 µL of FBS-free medium per well in a 12-well plate and starved at least 14h at 23°C. After starvation, either 10% (v/v) of FBS, 10% (v/v) of 1X PBS, or increasing concentrations of commercial Apolipoprotein B100 (#A5353-0.5MG, Sigma-Aldrich) in combination with 0.3 mM CaCl_2_ were added per condition. Cells were immediately put in 50 rpm orbital agitation at room temperature and monitored for 24h. Aggregates were imaged after 24h agitation at 10 distinct locations throughout each well at 10X magnification using transmitted light (brightfield) on an Eclipse TS100 Nikon epifluorescence inverted microscope equipped with an Intensilight C-HGFI Illuminator and a DS-Fi2 Camera Head. All experiments were performed in two biological replicates.

#### Whole LDLs dose response curve (related to Fig. 4C and Fig. S1D)

Adherent stage cells from a 2-day grown ∼90% confluence culture were scraped, homogenized and washed once with 15 mL of FBS-free media and allowed to starve overnight in a tube at 23°C. After starvation, 8*10^5^ cells were seeded in 160 µL of FBS-free media per well in a 96-well ultra-low attachment microplate and allowed to settle for 2 hours at room temperature. Whole LDLs (#L7914, Sigma-Aldrich) were first pre-washed from their storage buffer with Amicon Ultra 30 kDa MW cutoff centrifugal filters (#UFC903024, Millipore) and resuspended in 1X PBS buffer. To quantify the concentration of LDLs, the absorbance at 280 nm was measured with a BioTek Syngergy H1 plate reader. This value was used to calculate the concentration (µg/mL) of protein content. Since LDLs are ∼25% protein (75), this value was multiplied by 4 to yield the concentration (µg/mL) of entire LDL content. To induce aggregation, 60 µL containing 0-400 µg/mL of washed LDLs were added in each well in a 1:3 serial dilution (diluted in FBS-free media). Aggregates were imaged 90 min after induction at 5X magnification using transmitted light (brightfield) in a Leica DMi1 inverted microscope. All experiments were performed in three biological replicates.

#### Whole FBS vs. LDL-deficient FBS dose response curves (related to Fig. 4D, Fig. S1D and Fig. S10D)

Adherent stage cells from a 2-day grown ∼90% confluence culture were scraped, homogenized and washed once with 15 mL of FBS-free media and allowed to starve overnight in a tube at 23°C. After starvation, 8*10^5^ cells were seeded in 160 µL of FBS-free media per well in a 96-well ultra-low attachment microplate and allowed to settle for 2 hours at room temperature. To induce aggregation, 60 µL containing 0-30% (v/v) of whole FBS (#MT35011CV, Corning) or lipoprotein-deficient FBS (#S5394, Sigma-Aldrich) were added in each well in a 1:3 serial dilution (diluted in FBS-free media). Aggregates were imaged 90 min after induction at 5X magnification using transmitted light (brightfield) in a Leica DMi1 inverted microscope. All experiments were performed in three biological replicates.

#### qWestern Blots of FBS and LDL-deficient FBS (related to Fig. S10A-B)

An equivalent volume of ∼150 µg FBS was first diluted 1:2 with loading buffer (consisting of 2X Laemmli Sample Buffer (#1610737, Bio-Rad) plus a 5% (v/v) of 2-Mercaptoethanol (#M7154-25ML, Sigma-Aldrich)), further serially diluted 1:2 in loading buffer and boiled for 15 min at 99°C while mixed at 700 rpm. The same procedure (using the same starting volume) was followed to prepare LDL-deficient FBS samples. The same volume of all samples (corresponding to ∼15 µg starting FBS dilution) was separated by SDS-PAGE using 4-20% polyacrylamide Mini-Protean® TGX^TM^ Gels (#456-1094, Bio-Rad) and wet-transferred to Nitrocellulose membranes (#1620112, Bio-Rad) overnight at 4°C in 1X Tris-Glycine SDS buffer (#T7777-1L, Sigma-Aldrich) containing 20% (v/v) Methanol (#1.06018.2500, Supelco). Membranes were then blocked with 5% (w/v) skimmed milk in 1X PBS with 0.1% (v/v) Tween20 (#P1379-500ML, Sigma-Aldrich), hereafter PBS-T, during 1 hour at room temperature. Proteins were probed with 1:2000 chicken anti-LDL primary antibody (#GW20089F, Sigma-Aldrich) diluted in PBS-T overnight at 4°C and detected with 1:2000 HRP-conjugated goat anti-chicken secondary antibody (#A16054, Invitrogen) diluted in PBS-T during 1 hour at room temperature. Samples were finally developed in a 1:1 mix of Peroxide Solution and Luminol from the SuperSignal^®^ West Pico Chemiluminescent Substrate (#34078, ThermoFisherScientific) and visualized in a ChemiDoc^TM^ Touch Gel Imaging System Transiluminator (#1708370, Bio-Rad). Serial dilutions of commercial LDL (#SAE0053, Sigma-Aldrich) were prepared, separated and detected as before as a control for LDL detection using 1:2000 chicken anti-LDL primary antibody.

Western blot images were analyzed using Fiji Imaging Software version 2.3.0/1.53f (74). Images were first converted to 8-bit gray-scale. Then, the “Subtract Background” command from the “Process” menu was used to remove image background, setting the rolling ball radius parameter to 50 pixels. Next, a rectangular section covering high molecular weight bands in the first lane was drawn using the rectangular selection tool from the Fiji toolbar. The first lane was set using the “Select First Lane” command from the “Gels” option in the “Analyze” menu. This step was repeated using the “Select Next Lane” command in the same path. After setting rectangle sections in all lanes, the “Plot Lanes” command was used to draw a profile plot of each lane representing the relative density of the contents of the rectangle over each lane. Next, the wand tool from the Fiji toolbar was used to select each lane’s plot, and its corresponding measurements were recorded in the “Results” window. Finally, the “Label Peaks” command in the same path was used to label each lane’s plot with its size, expressed as a percentage of the total size of all of the highlighted lane plots. The percentages values of the resulting profile plots in each lane were normalized relative to values from the first FBS sample dilution.

### Lipids are the active components of low-density lipoproteins

#### Protein Free LDL-like particles (pfLDLs) dose response curve (related to Fig. 4E, Fig. S11 and Table S2)

Adherent stage cells from a 2-day grown ∼90% confluence culture were scraped, homogenized and washed once with 15 mL of FBS-free media and allowed to starve overnight in a tube at 23°C. After starvation, 8*10^5^ cells were seeded in 180 µL of FBS-free media per well in a 96-well ultra-low attachment microplate and allowed to settle for 2 hours at room temperature. Protein free LDL-like particles (pfLDLs) were prepared according to Vauhkonen *et al.,* (59) by first mixing POPC (#850457C, Avanti Polar Lipids), 16:0 SM (#860584C, Avanti Polar Lipids), cholesterol (#C8667, Sigma-Aldrich), triolein (#T7140, Sigma-Aldrich), and cholesteryl oleate (#C9253, Sigma-Aldrich) in a 35:12:40:21:100 mol/mol lipid mixture. The chloroform storage solvent was evaporated under gentle stream of nitrogen and then the dried lipid mixture was dissolved in isopropanol (20 µmol of phospholipid/mL). The solution (corresponding to 200 nmol of phospholipid) was heated to 50°C in an oven, drawn into a heated microsyringe and injected into 2 mL of rapidly stirred 1X PBS buffer at 10°C. The size and distribution of particles was confirmed using a DLS Zetasizer Nano and JEOL JEM 1010 TEM and the composition of particles was verified by LCMS using Waters Synapt G2S QTOF before testing. The pfLDLs were first pre-washed with Amicon Ultra 30 kDa MW cutoff centrifugal filters (#UFC903024, Millipore) and resuspended in 1X PBS buffer. Next, to induce aggregation, 20 µL of washed pfLDLs were added in each well in a 1:2 serial dilution (diluted in 1x PBS buffer) to a final concentration ranging from 0-153 µg/mL. Aggregates were imaged 90 minutes after induction at 5X magnification using transmitted light (brightfield) in a Leica DMi1 inverted microscope. All experiments were performed in triplicate.

### Dynamic lipoprotein concentration regulates the aggregative state of *Capsaspora*

#### LDL-depletion experiment (related to Fig. 5A, Fig. S4A-B and Fig. S12)

To assess LDL-depletion over time, aggregates were induced with 10% (v/v) FBS using orbital agitation and monitored during 80h (see *Re-aggregation experiment using orbital agitation, related to Fig. S3 and Fig. S1B assay)*. Around 1 mL of sample supernatant was collected for each timepoint during the aggregation-disaggregation process (timepoints 0-80h agitation). A new well was sampled for each timepoint. Samples were harvested at 5000xg for 5 min at room temperature and the supernatant was collected and stored at -20°C until further use. The same volume in all timepoint samples (equivalent to ∼30 µg of growth media) was diluted 1:2 with loading buffer and boiled for 15 min at 99°C while mixed at 700 rpm. Around 20 µg of control sample (and the corresponding equivalent volumes for the rest of the timepoints) were separated by SDS-PAGE and detected using 1:2000 chicken anti-LDL primary antibody as before. Western blot images were analyzed as before, normalizing the percentages values of the resulting profile plots in each lane relative to values from timepoint 0h.

#### Immunofluorescence Microscopy (related to Fig. 5B and Fig. S13)

Adherent cells from a ∼90% confluence culture were scraped, homogenized and counted to seed 7.5*10^5^ cells in 1 mL of growth media per well in a 12-well plate (#55428, Nunc/DDBioLab) and incubated overnight at 23°C. After incubation, cells were put at 50 rpm orbital agitation (Celltron Bench-Top Shaker, INFORS-HT) during 24h at room temperature to induce aggregate formation. For aggregate isolation, ∼300 µL of volume containing *Capsaspora* aggregates in suspension were pipette-picked using a 1000 µL pipette tip (around 1 cm of the tip was previously cut) and transferred to an Eppendorf tube. Next, 8% formaldehyde (#F8775-25ML, Sigma-Aldrich) prediluted in 1X PBS were added to the tube, reaching a final 4% formaldehyde concentration, and immediately put for 15 min at 50 rpm orbital agitation at room temperature. After incubation, the tube was removed from agitation and incubated an extra 15 min to let aggregates sink (reaching a total fixation time of 30 min at room temperature). Next, aggregates were washed twice by removing around 90% of the volume of the tube and replaced with 1 mL 1X PBS, letting aggregates sink as before. After washing, aggregates were blocked with Blocking Solution (1% BSA (#A3294-10G, Sigma-Aldrich) and 0.1% Triton X-100 (#X100, Sigma-Aldrich) pre-diluted in 1X PBS) for 1h at room temperature. Aggregates were then incubated with 1:100 chicken anti-LDL primary antibody (#GW20089F, Sigma-Aldrich) overnight at 4°C. After incubation, aggregates were washed twice as before with 500 µL Blocking Solution. Then, aggregates were incubated with 1:2000 of goat anti-chicken Alexa Fluor 488 secondary antibody (#H11039, Invitrogen) for 30 min at 50 rpm orbital agitation at room temperature. Next, a 1:100 of Alexa Fluor^TM^ 350 Phalloidin (#A22281, ThermoFisherScientific) was added and samples were incubated for extra 30 min at 50 rpm orbital agitation at room temperature. After incubation, the tubes were placed in the tube rack for 15 min to let aggregates sink. Aggregates were washed twice as before with 500 µL 1X PBS. Aggregates were then resuspended in ∼30 µL of 1:100 DRAQ5 (#62251, ThermoFisherScientific) pre-diluted in 1X PBS. Around 10-20 µL of each sample were added to a cover slide (previously washed with distilled water and 70% ETOH and later treated with 10 µL of Poly-L-Lysine (#P4832, Sigma-Aldrich)) and air-dried. Samples were mounted with ProLong Gold antifade reagent (#P36930, ThermoFisherScientific) and observed at 40X magnification on a Leica TCS SP5 II inverted confocal microscope. Acquisition settings were adjusted using a negative control without primary antibody.

A control of LDL incorporation in adherent cells **(Fig. S13B)** grown in growth medium (containing 10% (v/v) FBS) was prepared as follows. Adherent cells from an 80-90% confluent culture were scraped and collected by harvesting at 5000xg for 5 min at room temperature. Cells were then washed twice with 10 mL 1X PBS, centrifuging as before. Next, cell pellets were gently resuspended by pipetting up and down with 750 µL 1X PBS. An extra 750 µL of 8% formaldehyde (pre-diluted in 1X PBS) were added to reach a final 4% formaldehyde, and cells were fixed for 7 min at room temperature in a rotating shaker. After incubation, cells were washed twice with 1.5 mL 1X PBS by gently pipetting up and down, incubating the tube in a rotating shaker for 2 min, and by harvesting cells at 5000xg for 90 seconds at room temperature. Next, cells were blocked in 800 µL of Blocking Solution for 1h at room temperature in a rotating shaker. After blocking, a 1:100 chicken anti-LDL primary antibody was directly added and incubated overnight at 4°C in a rotating shaker. Then, cells were washed twice in 500 µL Blocking Solution as before and incubated 30 min at room temperature on a rotating shaker with 1:2000 of goat anti-chicken Alexa Fluor 488 secondary antibody. A 1:100 of Texas Red^TM^-X Phalloidin (#T7471, ThermoFisherScientific) was directly added and incubated for an extra 30 min at room temperature in a rotating shaker. Next, cells were washed twice with 500 µL 1X PBS and resuspended in ∼20 µL 1:100 DRAQ5 pre-diluted in 1X PBS. Samples were then added to a cover slide as before, air-dried and mounted with ProLong Gold antifade reagent. Adherent cells were observed at 63X magnification using immersion oil on a Zeiss Axio Observer Z.1 epifluorescence inverted microscope equipped with LED illumination and an Axiocam 503 mono camera. Acquisition settings were adjusted using a negative control without primary antibody.

### Only viable *Capsaspora* cells aggregate

#### Aggregation induction in formalinized cells (related to Fig. 5C and Fig. S1B)

Adherent cells from a ∼90% confluence culture were scraped, homogenized and harvested at 5000xg during 5 min at room temperature. Cells were then washed twice with 10 mL of filter-sterilized 1X PBS (#P5368-10 PAK, Sigma-Aldrich) containing 0.3 mM CaCl_2_ (#C1016, Sigma-Aldrich) and 0.3 mM MgCl_2_ (#M8266, Sigma-Aldrich). After the second wash, cells were resuspended in 500 µL of the previous buffer solution and homogenized by pipetting up and down to prevent random cell clump fixation. Next, an additional 500 µL containing 0-32% of increasing concentrations of formaldehyde (#F8775-4X25ML, Sigma-Aldrich) diluted in the previous buffer solution were added for each condition. Samples were homogenized by gently pipetting and fixed for 10 min at room temperature on a rotating shaker. After fixation, cells were harvested at 6500xg during 5 min at room temperature and washed with 1 mL of the previous buffer solution. Cells were later resuspended to an appropriate volume of FBS-free medium to reach a cell concentration of 8.33*10^6^ cells/mL. Next, 7.5*10^5^ cells were seeded in 900 µL of FBS-free medium per well in a 12-well plate and starved at least 14h at 23°C. After starvation, either 10% (v/v) of 1X PBS or 10% (v/v) of FBS were added per condition. Cells were immediately put in 50 rpm orbital agitation at room temperature and monitored for 24h. Aggregates were imaged after 2-4h agitation at 10 distinct locations throughout each well at 10X magnification using transmitted light (brightfield) on an Eclipse TS100 Nikon epifluorescence inverted microscope equipped with an Intensilight C-HGFI Illuminator and a DS-Fi2 Camera Head. All experiments were performed in three biological replicates, testing each condition per duplicate.

## ACKNOWLEDGMENTS

We thank the Light Microscopy Center at Indiana University for support in image acquisition and analysis (funding provided by the NIH grant NIH1S10OD024988-01) and the Advanced Light Microscopy Unit of the CRG for support on images acquisition. We also thank the Indiana University Nanoscale Characterization Facility, Electron Microscopy Center, and Laboratory for Biological Mass Spectrometry for use of their instruments. We want to special thank Claudio Scazzocchio for fruitful discussions and Sebastián R. Najle for feedback and support in early work on *Capsaspora* aggregation. We acknowledge Omaya Dudin for feedback and support in image acquisition and analysis, Michelle M. Leger for feedback on the manuscript and Koryu Kin for support in immunocytochemistry experiments, image acquisition and analysis and stimulating discussions. We also thank many people from the lab for insights and support that helped advance this project.

This work was supported by a NIH grant (R35GM138376) to J.P.G. and grants BFU2017-90114-P from Ministerio de Economía y Competitividad (MINECO), Agencia Estatal de Investigación (AEI), and Fondo Europeo de Desarrollo Regional (FEDER) and PID2020-120609GB-I00 by MCIN/AEI/10.13039/501100011033 and “ERDF A way of making Europe” to I.R.-T. N.R.-R. was supported by a “Formación del Profesorado Universitario (FPU13/01840)” predoctoral fellowship from Ministerio de Educación, Cultura y Deporte (MECD) and R.Q.K. was supported by a NIH training grant (T32GM131994).

## SUPPLEMENTARY FIGURES

**Fig. S1.**
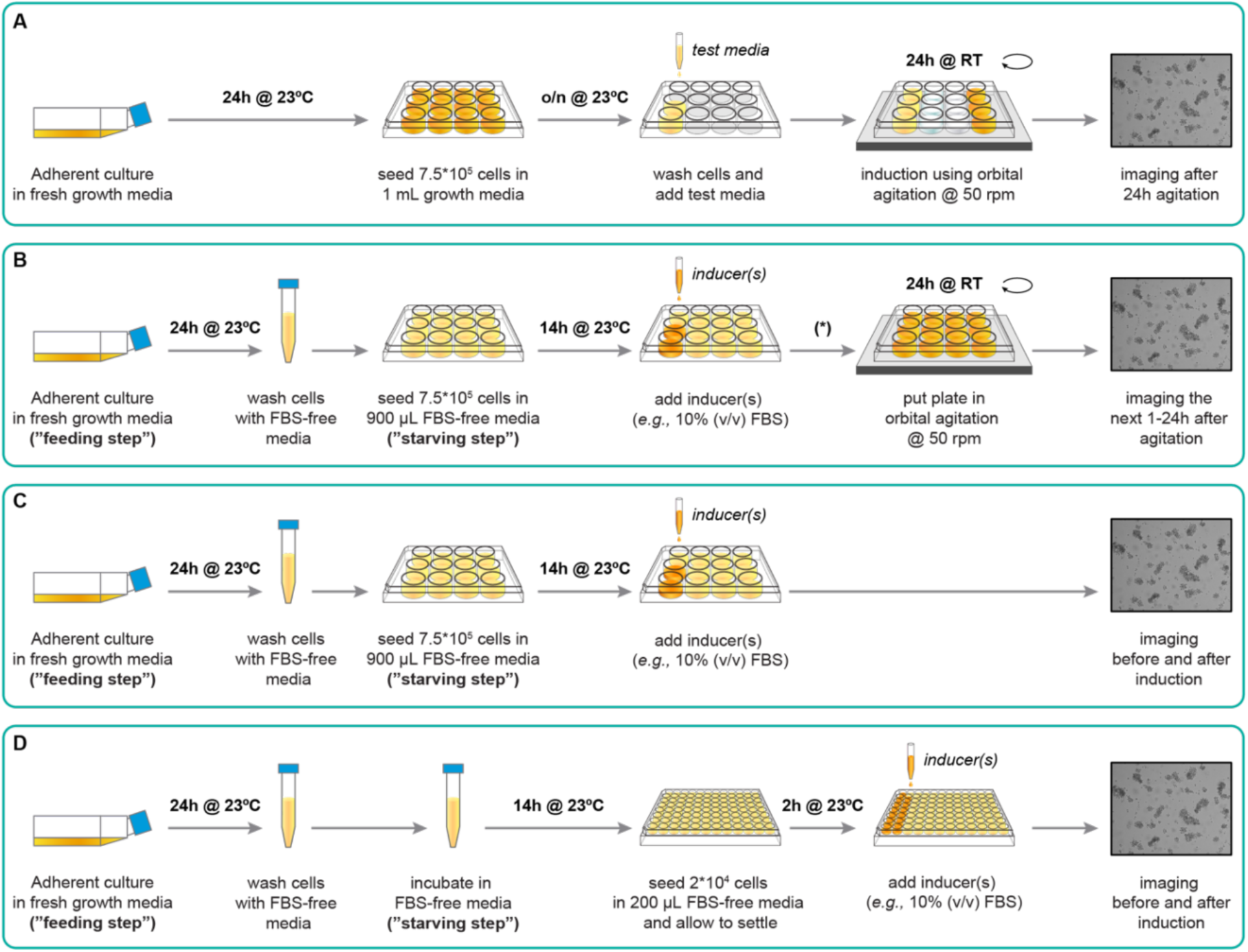
Experimental designs of *Capsaspora* aggregation assays induced by FBS and related components in the presence and absence of orbital agitation. (A) Aggregation assay in a 12-well plate induced by orbital agitation, modified from the original aggregation assay developed in (35). (B) Aggregation assay induced by FBS and related components using 12-well plates in orbital agitation, including a feeding-starving step to control for background aggregation caused by intracellular or extracellular remnants of FBS. (*) Note that after inducer(s) addition, *Capsaspora* cells start aggregating, even in the absence of orbital agitation. (C) Aggregation assay induced by FBS and related components using 12-well plates as in B in the absence of orbital agitation. (D) Aggregation assay induced by FBS and related components using ultra-low attachment 96-well plates in the absence of orbital agitation, including a feeding-starving step.

**Fig. S2.**
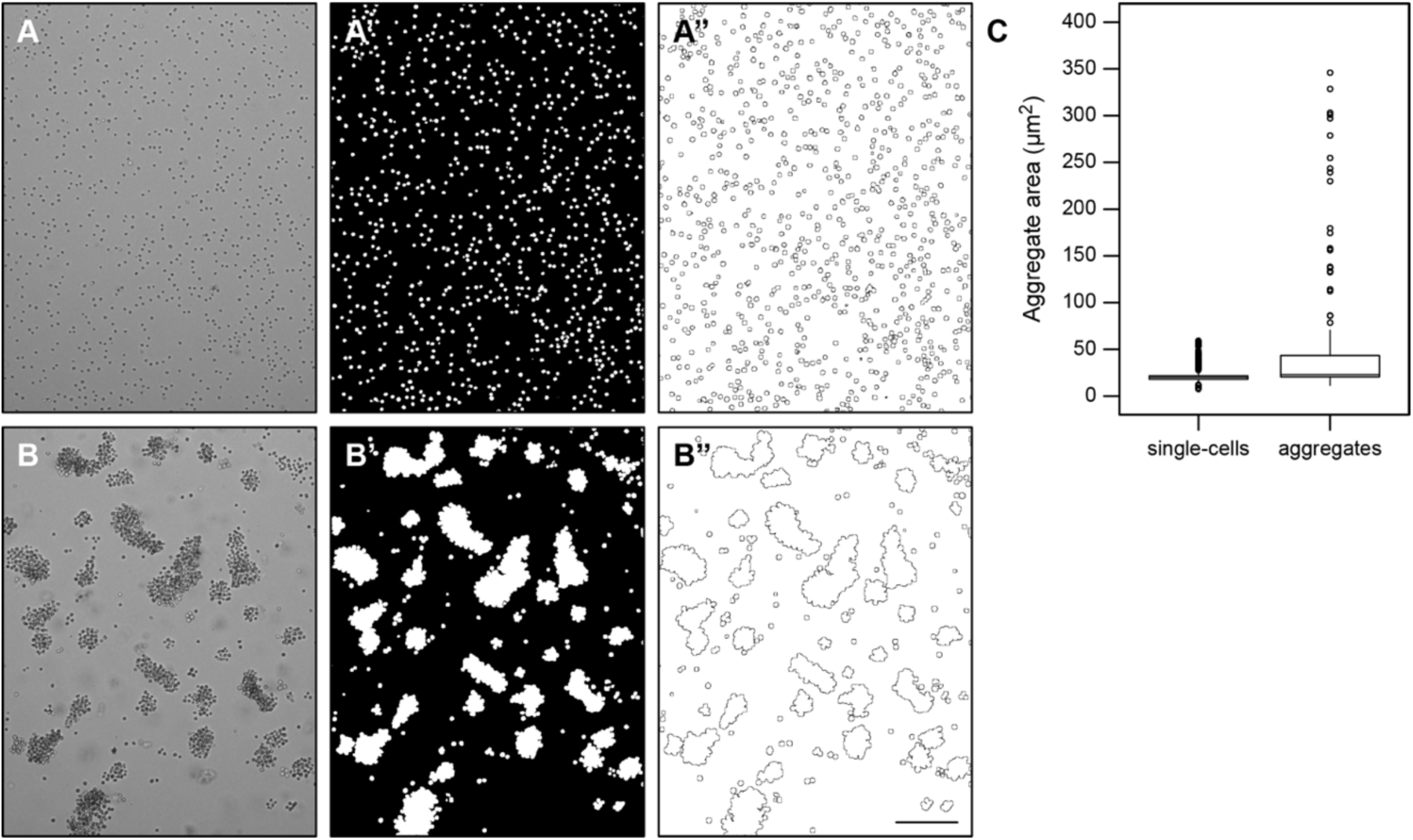
Automated image analysis distinguishes between *Capsaspora* aggregates and single cells. Automated image analysis using a batch processing macro script (74) transforms microscopy images of single-cells (A) and aggregates (B) into binary images (A’-B’) which are then outlined to measure particle sizes (A’’-B’’). (C) Distribution of particle area of single cells or aggregates quantified from images in (A’’-B’’). Scale bar represents 100 μm.

**Fig. S3.**
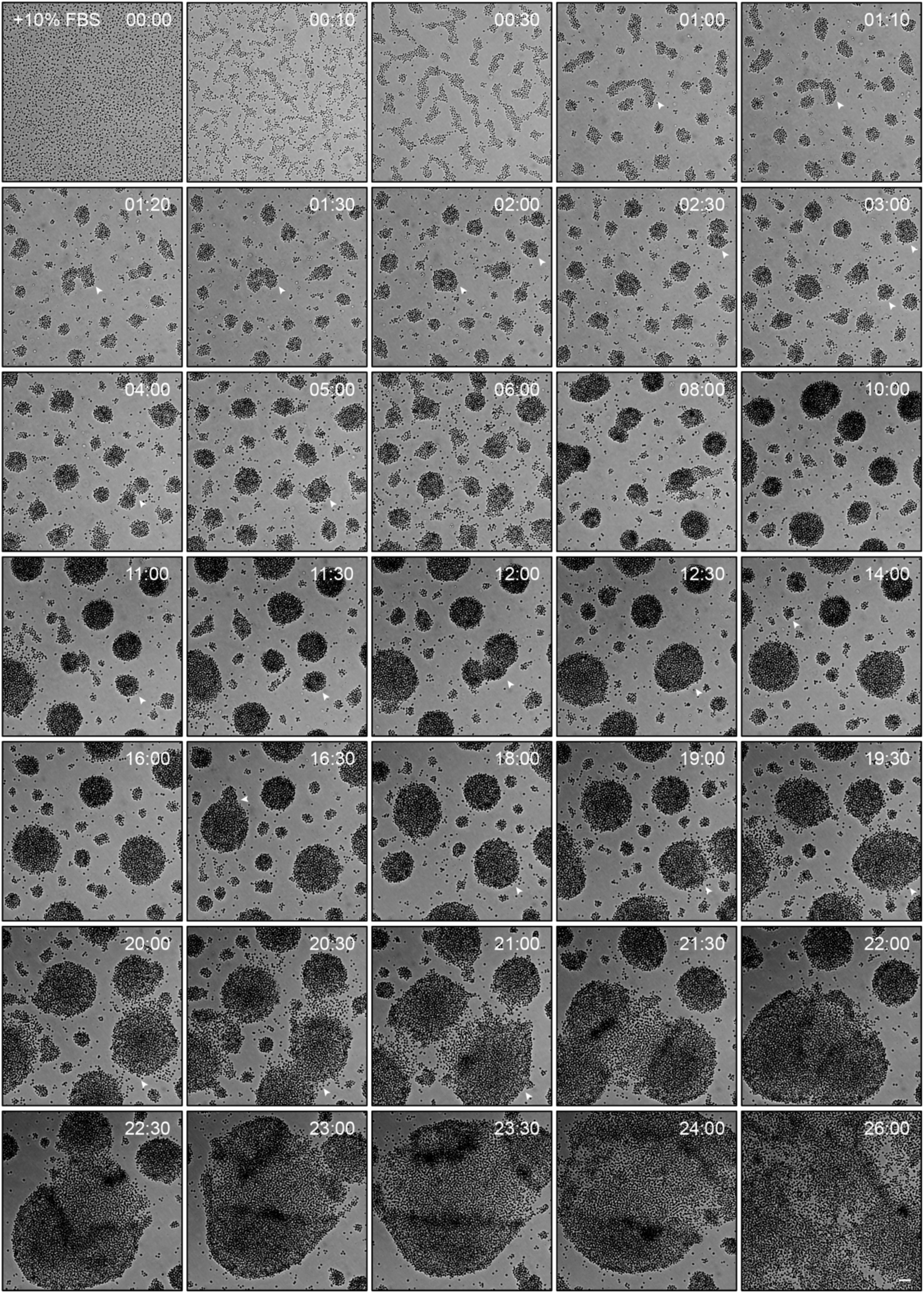
Chemical cues from FBS are sufficient to induce aggregation in the absence of physical agitation. Images of aggregate formation induced upon 10% (v/v) of FBS addition (time 0) in ultra-low attachment plates **(Fig. S1C assay)**. White arrowheads track early aggregate formation and aggregate fusion events (agglomeration). Time represents hh:mm. Scale bar represents 100 μM. Images related to **Movie 3**.

**Fig. S4.**
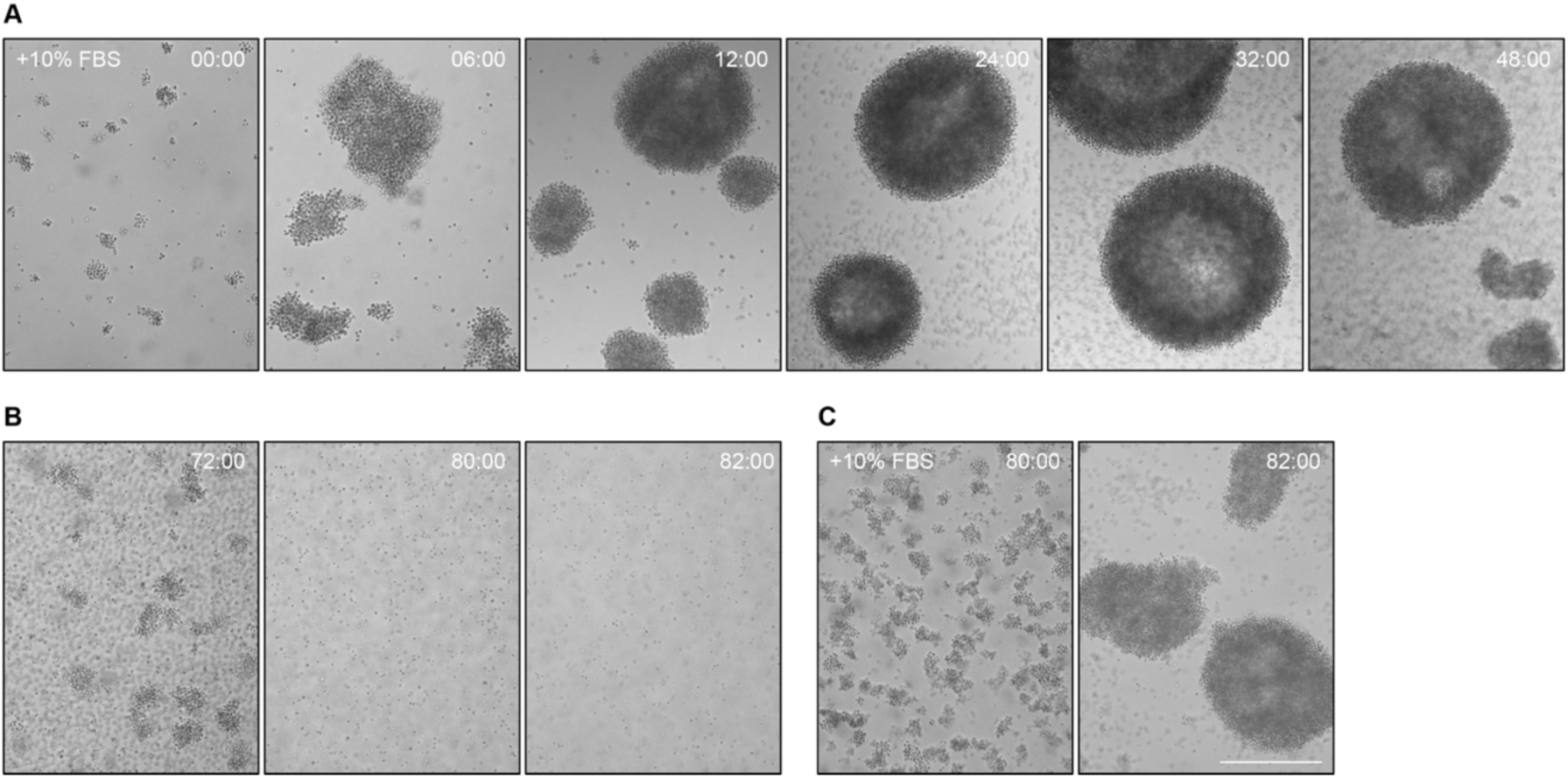
Re-addition of FBS in disaggregated cells induces re-aggregation in *Capsaspora*. (A) *Capsaspora* cells aggregate upon addition of 10% (v/v) FBS during 48h in agitation conditions **(Fig. S1B assay)**. (B) A few hours later, they disaggregate into single cells again (72-82h). (C) A 10% (v/v) FBS re-addition after 80h agitation in disaggregated cells induces re-aggregation (80h-82h). Time represents hh:mm. Scale bar represents 100 μm. Figure related to **Fig. 2G** & **4E**.

**Fig. S5.**
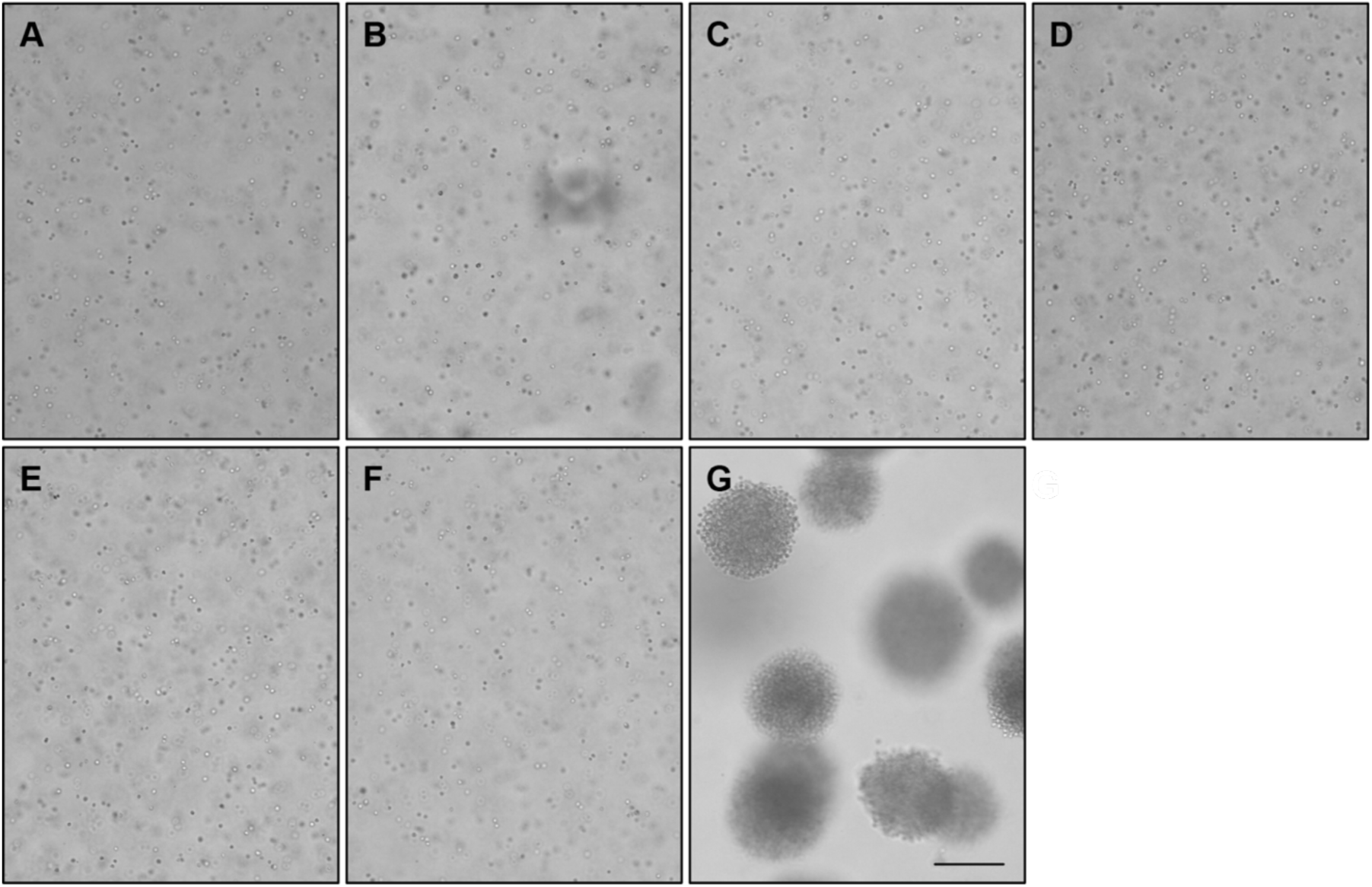
Insoluble salt(s) from <3 kDa FBS fraction induce aggregation when combined with >30 kDa FBS fraction in *Capsaspora*. The aggregation activity of <3 kDa FBS fraction components was tested by treating *Capsaspora* cells in serum-free media with >30 kDa FBS fraction (in all conditions) and either one of the following components: (A) nothing else added (negative control), (B) chloroform-soluble components, (C) 100% methanol-soluble components, (D) 80% methanol-soluble components, (E) 20% methanol-soluble components, (F) water-soluble components, (G) final insoluble components. Dark shadow in (B) was due to chloroform etching the polystyrene plate. Scale bar represents 100 μm. Figure related to **Fig. 3.**

**Fig. S6.**
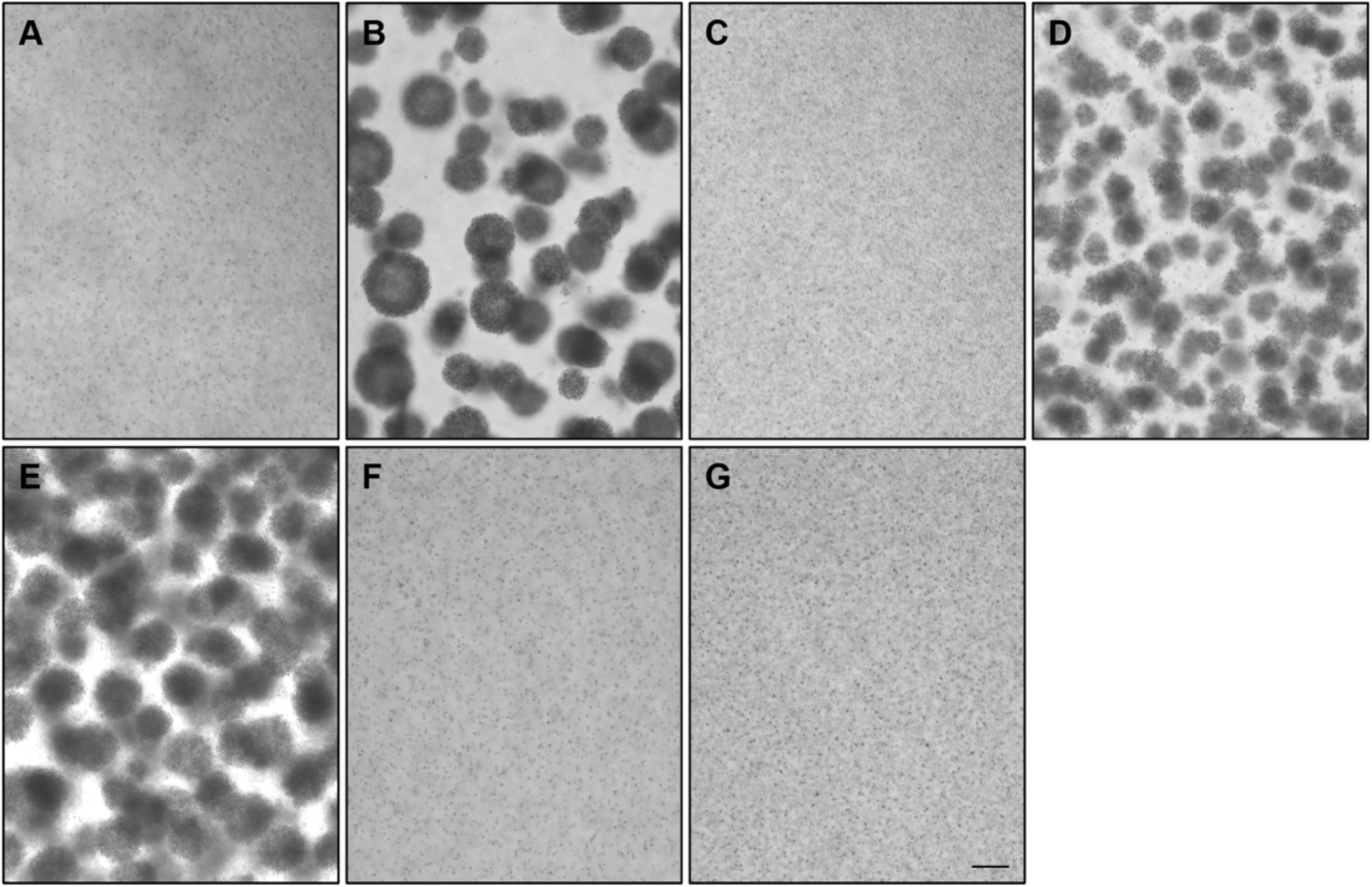
Ammonium sulfate-precipitated fractions induce aggregation in *Capsaspora*. The aggregation activity of >30 kDa FBS fraction components was tested by treating *Capsaspora* cells in FBS-free media with 0.2 mM CaCl_2_ (in all conditions) and either one of the following components: (A) nothing else added (negative control); (B) complete >30 kDa FBS components (positive control); (C) re-dissolved 20% ammonium sulfate precipitate; (D) re-dissolved 40% ammonium sulfate precipitate; (E) re-dissolved 65% ammonium sulfate precipitate; (F) re-dissolved 80% ammonium sulfate precipitate; and (G) washed supernatant remaining from 80% ammonium sulfate precipitation. Scale bar represents 100 μm.

**Fig. S7.**
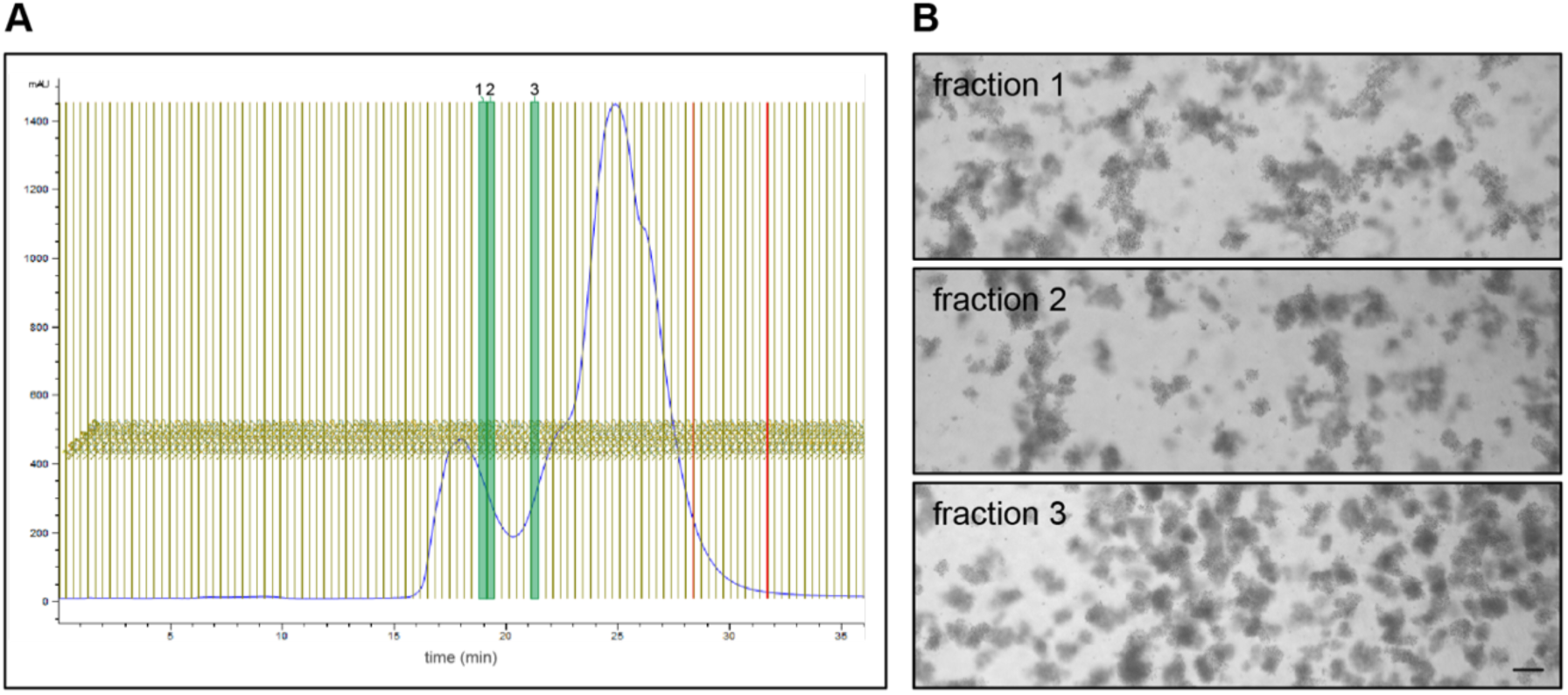
Anion exchange chromatography fractions induce aggregation in *Capsaspora*. (A) Chromatogram (254 nm absorbance) of anion exchange chromatography of active >30 kDa FBS ammonium sulfate precipitate fractions. The fractions that exhibited the most robust aggregation activity are highlighted in green. (B) Microscopy images of active fractions 1-3 from chromatogram in A inducing aggregation. Scale bar represents 100 μm.

**Fig. S8.**
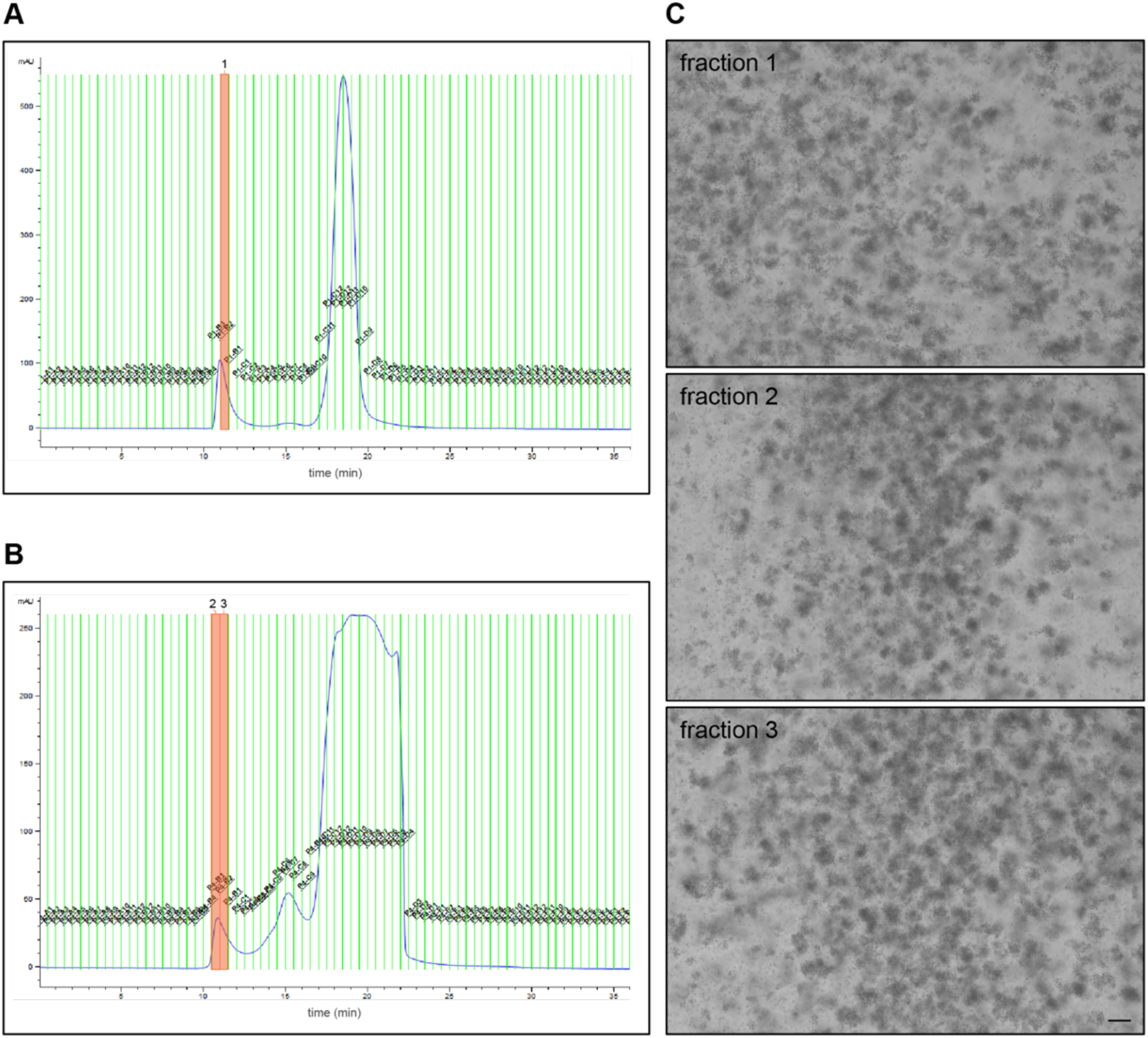
Size exclusion chromatography fractions induce aggregation in *Capsaspora*. Chromatograms (254 nm absorbance) of size exclusion chromatography of the active >30 kDa FBS anion exchange fractions resulting from **Fig. S7B**. Chromatograms resulted from (A) pooled fractions “1” and “2”, and (B) fraction “3” (related to **Fig. S7B**). The fractions that exhibited the most robust aggregation activity are highlighted in orange. These fractions eluted very early (within the first ∼12 minutes), indicating a large size for the active FBS component. (C) Microscopy images of active fractions from chromatograms A-B inducing aggregation. Scale bar represents 100 μm.

**Fig. S9.**
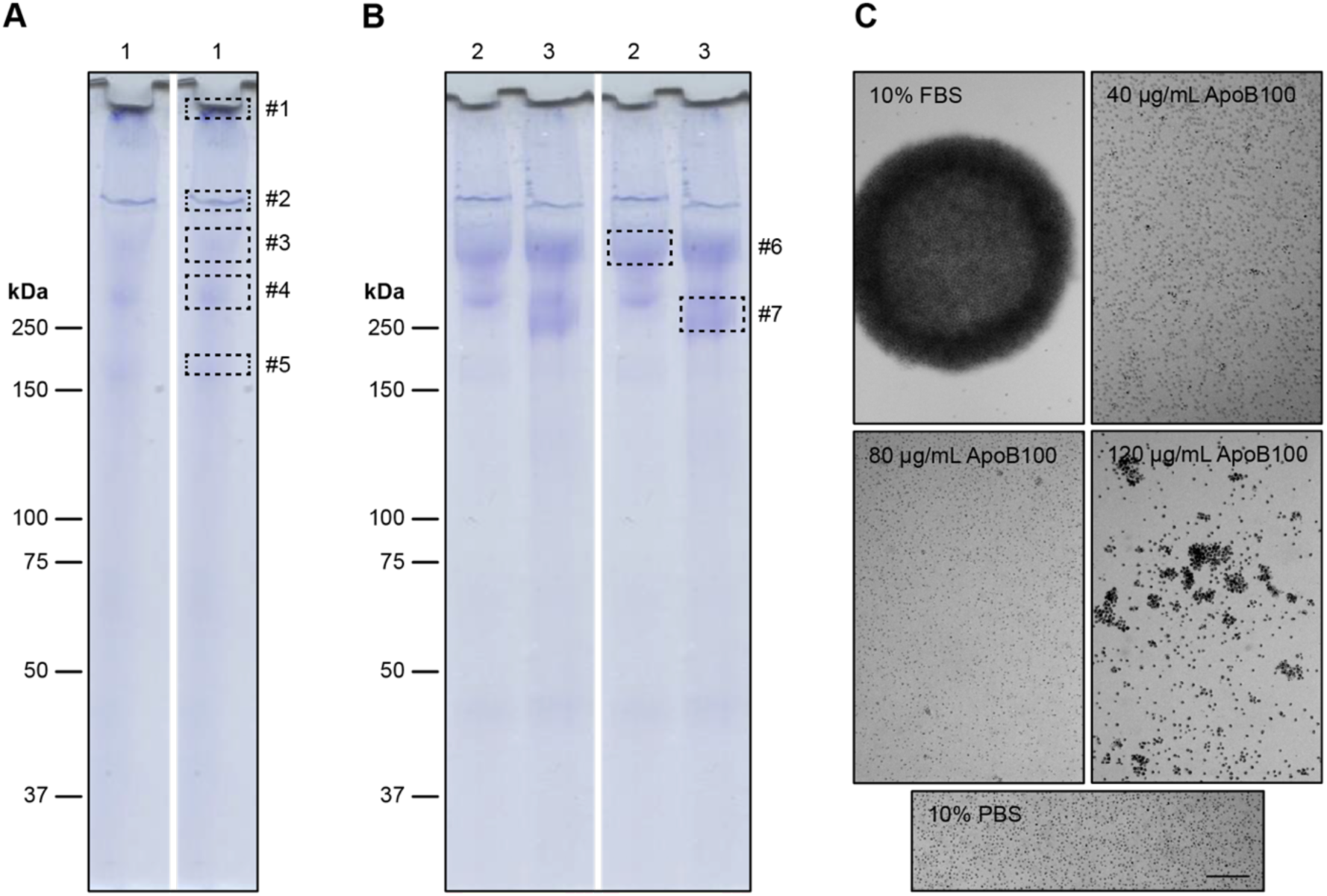
Major SDS-PAGE bands in active size exclusion fractions from >30 kDa FBS fraction are Apolipoprotein B100. (A-B) Coomassie-stained gels of active fractions “1” (shown in A) and “2” and “3” (shown in B) from size exclusion chromatography, related to supplementary **Fig. S8**. Right images in A-B show boxes around the bands (#1-7) that were excised and subjected to in-gel trypsin digestion and LC-MS/MS. The primary protein of each band was Apolipoprotein B100 (ApoB100, see **Table S1**). (C) ApoB100 in combination with 0.3 mM CaCl_2_ does not recapitulate the aggregation activity of 10% (v/v) FBS. At 120 µg/mL, ApoB100 appears toxic. A 10% (v/v) of 1X PBS was used as a negative control. Scale bar represents 100 μm. Microscopy images representative of each test condition in **Fig. 4B**.

**Fig. S10.**
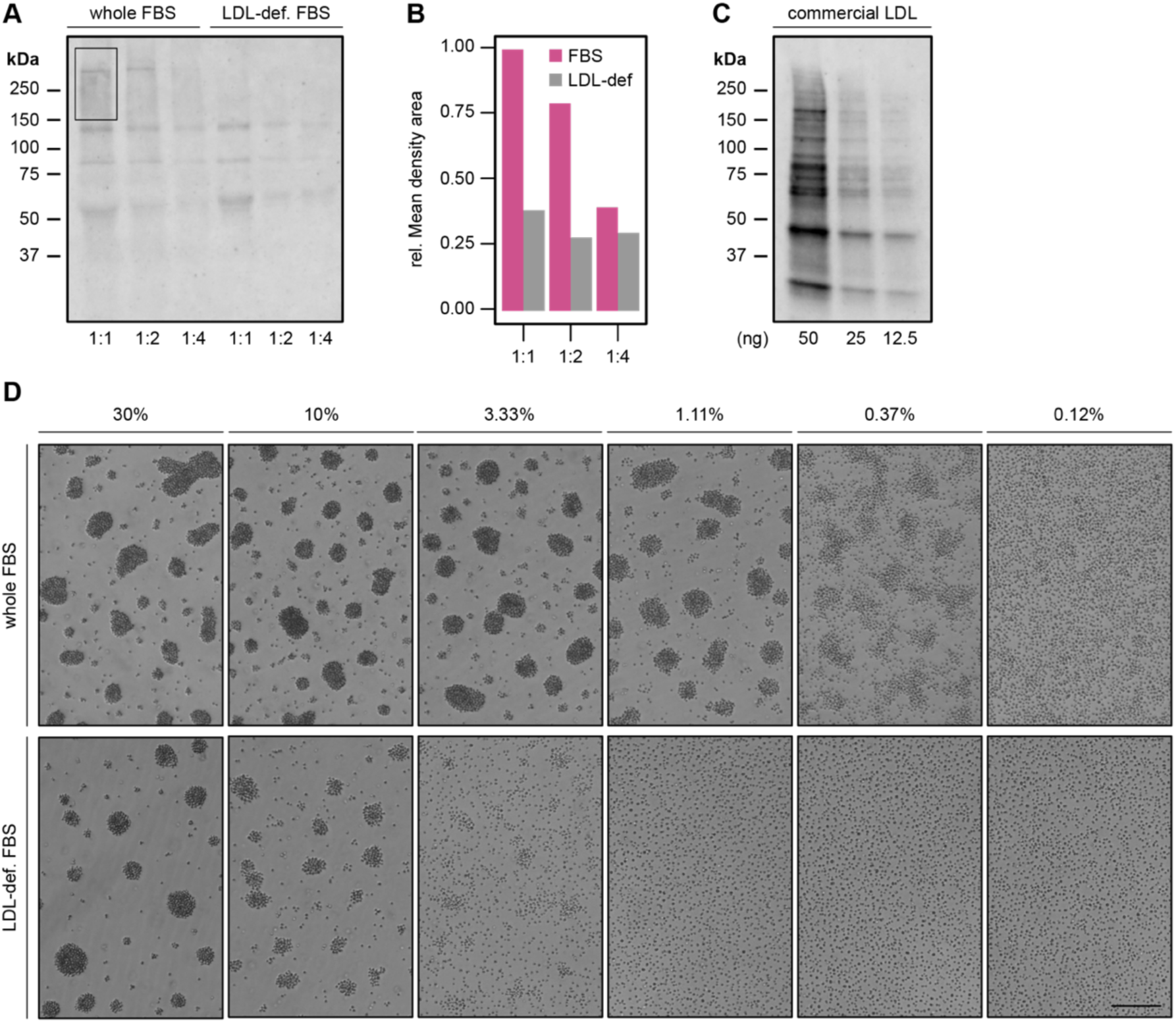
LDL-deficient FBS has reduced aggregation activity. (A) Western Blot analyses probed with anti-LDL antibody against 1:2 serial dilutions of whole FBS (starting from ∼15 μg of whole FBS [1:1]) and an equivalent volume of LDL-deficient FBS show that LDL-deficient FBS has significantly less LDL content. (B) Mean density areas relative to whole FBS derived from A. (C) Control for LDLs detection probed using anti-LDL antibody against 1:2 serial dilutions of commercial LDL. (D) Microscopy images of aggregates induced with 1:3 serial dilutions (expressed as % (v/v)) of whole FBS versus LDL-deficient FBS. Scale bar represents 100 μm. Figure related to **Fig. 4D**.

**Fig. S11.**
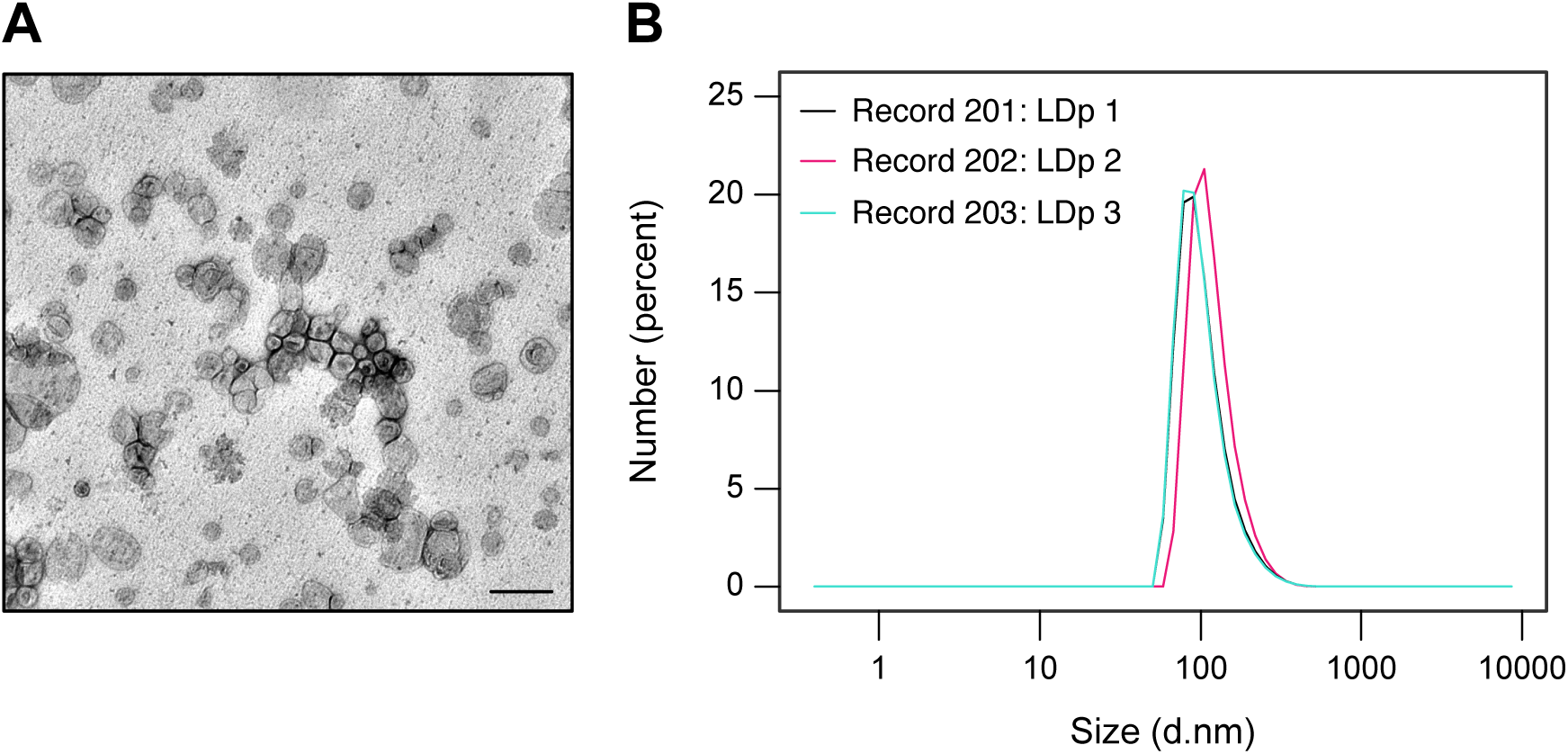
Formation of protein-free LDL particles (pfLDLs). (A) Transmission electron micrograph (TEM) of protein-free LDL particles. Scale bar: 0.2 µm. (B) Dynamic light scattering (DLS) measurements of the distribution of particle diameters in nanometers (d. nm). The three curves are three individual measurements of the sample of assembled particles. The TEM and DLS measurements both indicate the particles were ∼100 nm in diameter. Figure related to **Fig. 4E**.

**Fig. S12.**
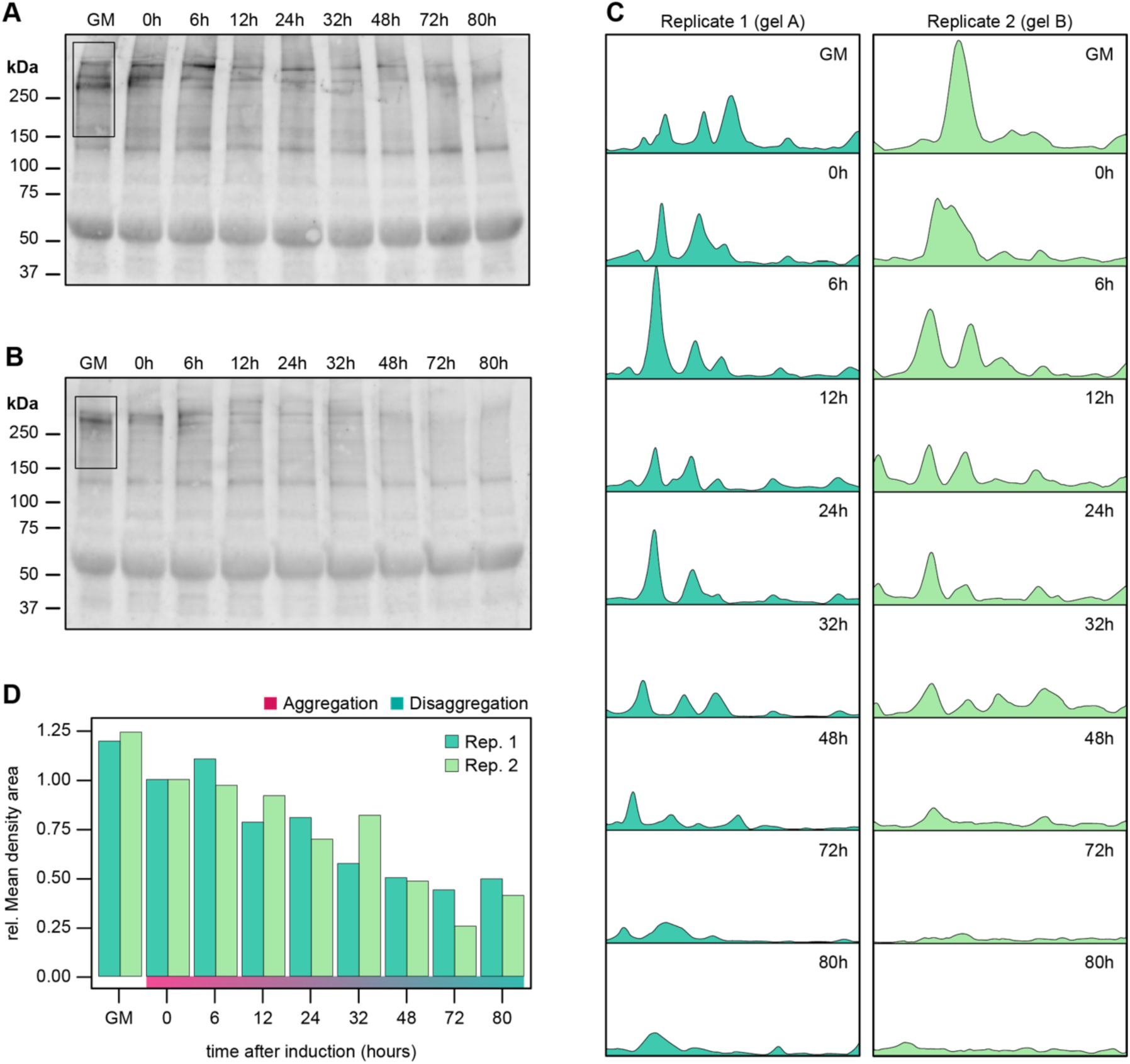
LDLs are depleted over time during the aggregation-disaggregation process. (A-B) Western Blot analyses probed with anti-LDL antibody of media supernatants collected from cells induced with 10% (v/v) FBS (0h) show major bands during aggregation (6-48h) and their progressive disappearance during disaggregation (72-80h). An equivalent volume to ∼30 μg of growth media (GM) was taken for all samples in two biological replicates (gel A, replicate 1; gel B, replicate 2). (C) Profile plots representing the density of the band contents in each lane (area inside rectangle) from gel images in A-B. Peaks correspond to dark bands in the original gel image as follows: higher peaks represent darker bands and wider peaks represent bands that cover a wider size range on the original gel. (D) Mean density areas relative to 0h timepoint derived from C. Figure related to **Fig. 5A** & **S4A-B**.

**Figure S13.**
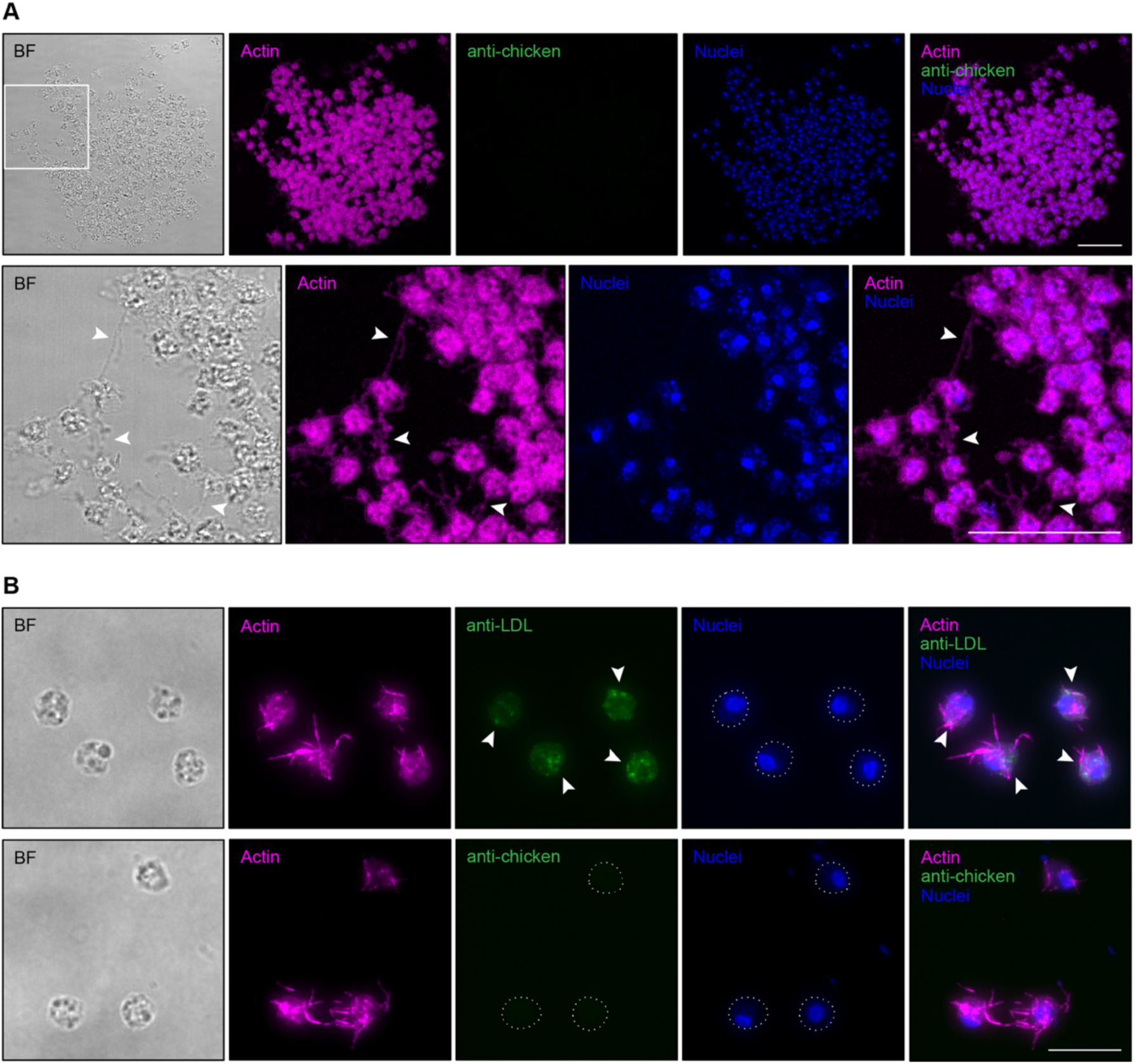
LDLs are incorporated in small foci inside the cytoplasm of *Capsaspora* cells. (A) Maximum z-projections of an aggregate stained with phalloidin to mark filamentous actin (magenta), including cell body and filopodia, an anti-chicken secondary antibody (green), and DRAQ5 to mark nuclei (blue). A representative brightfield (BF) stack image is shown. Lower panel represents a zoom-in of the aggregate (white square), showing cell-cell contacts through filopodia (white arrows). Scale bars represent 25 μm. Image corresponds to negative control without primary antibody related to **Fig. 5B**. (B) Adherent cells cultured in growth medium (containing 10% (v/v) FBS) also show intracellular reservoirs of LDLs (white arrows), but no signal along filopodia. Shown are representative maximum z-projections of cells stained as before (lower panel), or cells stained with an anti-LDL primary antibody (green, upper panel), showing intracellular foci of LDLs. Dotted lines indicate cell bodies. Scale bar represents 10 μm.

## SUPPLEMENTARY MOVIES LEGENDS

**Movie S1 (separate file). FBS induces *Capsaspora* aggregation and does not require orbital agitation for its activity.** Aggregation was induced by the addition of FBS (time 0) to a final concentration of 10% (v/v) after 9 min of imaging (left panel) without orbital agitation in 12-well plates **(Fig. S1C assay)**. Under these conditions, cells start forming aggregates after 21 min of induction, and continue aggregating during approximately 4h, after which the aggregates disintegrate into single-cells. A 10% (v/v) of 1X PBS was added as negative control (right panel). Aggregates were imaged every 3 minutes for 22 hours, and converted to a movie at 7 frames per second (fps). Time represents hh:mm. Scale bar represents 100 µm. Movie related to **Fig. 2C**.

**Movie S2 (separate file). Higher FBS concentration has a greater effect on *Capsaspora* aggregation in the absence of orbital agitation.** Aggregation was induced by the addition of FBS (time 0) to a final concentration of 30% (v/v) after 9 min of imaging (left panel) without orbital agitation in 12-well plates **(Fig. S1C assay)**. Under these conditions, cells actively aggregate during approximately 8h, after which the aggregates disintegrate into single-cells. A 30% (v/v) of 1X PBS was added as negative control (right panel). Aggregates were imaged every 3 minutes for 22 hours, and converted to a movie at 7 fps. Time represents hh:mm. Scale bar represents 100 µm. Movie related to **Fig. 2C**.

**Movie S3 (separate file). *Capsaspora* aggregates can be chemically induced and maintained for longer time in ultra-low attachment surfaces.** Aggregation was induced by the addition of FBS (time 0) to a final concentration of 10% (v/v) after 60 min of imaging (left panel) without orbital agitation in ultra-low attachment plates **(Fig. S1D assay)**. Under these conditions, cells immediately begin to aggregate upon induction. After 24h, the aggregates disintegrate into single-cells. Note that aggregates get larger over time, either by incorporating single cells (early stages of aggregation) or via smaller aggregates merging into larger aggregates (agglomeration). A 10% (v/v) of 1X PBS was added as negative control (right panel). Aggregates were imaged every 10 minutes for 42 hours, and converted to a movie at 7 fps. Time represents hh:mm. Scale bar represents 100 µm. Movie related to **Fig. 2E-F** and **Fig. S3**.

**Movie S4 (separate file). Dose-response of FBS shows distinct aggregation effect in the absence of physical agitation.** Aggregation was induced by the addition of FBS (time 0) to a final concentration of (A) 10% (v/v), (B) 5% (v/v), (C) 1.5% (v/v), and (D) 0.5% (v/v) after 60 min of imaging without orbital agitation in ultra-low attachment plates **(Fig. S1D assay)**. Under these conditions, cells immediately aggregate upon induction during approximately 24h, after which aggregates disintegrate into single-cells. Lower FBS concentrations (D) weakly induce aggregation. A 10% (v/v) of 1X PBS was added as negative control (E). Aggregates were imaged every 10 minutes for 42 hours, and converted to a movie at 7 fps. Time represents hh:mm. Scale bar represents 100 µm. Movie related to **Fig. 2E**.

**Movie S5 (separate file). Re-addition of FBS in disaggregated *Capsaspora* cells induces re-aggregation.** Aggregation was induced by the addition of FBS (time 0) to a final concentration of 10% (v/v) after 9 min of imaging without orbital agitation in 12-well plates **(Fig. S1C assay)**. Under these conditions, cells actively aggregate during approximately 7h, after which the aggregates disintegrate into single-cells. At this point (∼19h after induction), a 30% (v/v) of FBS was re-added to induce aggregation again (left panel). A 30% (v/v) of 1X PBS was re-added as a negative control (right panel). Aggregates were imaged every 3 minutes for 25 hours, and converted to a movie at 7 fps. Time represents hh:mm. Scale bar represents 100 µm. Movie related to **Fig. 2G**.

## SUPPLEMENTARY MATERIAL LEGEND

**Table S1 (separate file). Determination of protein content of active size exclusion fractions via mass spectrometry.** LC-MS/MS analysis of tryptic digests of all visible unique SDS-PAGE bands in active size exclusion fractions from >30 kDa FBS fraction **(Fig. S9A-B)**. The major protein in all 7 bands was Apolipoprotein B100 (ApoB100). Data in file <u>Table_S1_Compiled_MSMS_data.xlsx</u>.

**Table S2 (separate file). Relative composition of lipids in protein-free LDL particles.** LCMS data quantifying the relative composition of lipids in the protein-free LDL particles. Composition was calculated by comparing LCMS peak intensities in the assembled particles with LCMS peak intensities in a defined mixture of the lipids. Data in file <u>Table_S2_composition_of_lipid_particles.xlsx</u>.

